# Parallel CRISPR screens reveal pathways controlling the cell surface levels of the attractant receptor FPR1

**DOI:** 10.1101/2025.04.21.649864

**Authors:** Emel Akdoğan, Stefan M. Lundgren, Roarke A. Kamber, Michael C. Bassik, Sean R. Collins

## Abstract

Levels of receptors on the cell surface control the sensitivity of immune cells, which is central to achieving effective responses while avoiding inflammatory diseases. The chemoattractant receptor FPR1 activates strong chemotactic and cytotoxic responses in neutrophils. While its precise regulation is critical for appropriate responses, the mechanisms controlling its cell surface levels remain unclear. Here, we investigated the roles of both classic and unknown regulators. First, we found that multiple G protein-coupled receptor kinases (GRK2, GRK3, and GRK6), and both β-arrestin1 and 2 are important for FPR1 internalization. However, FPR1 uses multiple endocytic pathways, as cells lacking β-arrestins have strong, but incomplete defects in FPR1 internalization. Moreover, we performed two parallel genome-wide CRISPR/Cas9 screens for the regulators of FPR1 surface expression and internalization in a neutrophil-like cell line. These screens identified regulators of FPR1 surface expression, recycling, and endocytosis. We identified the formin mDia1 and the small GTPase Arf6 as specific regulators of FPR1 internalization, and we confirmed these phenotypes using chemical inhibitors in primary human neutrophils. Furthermore, our data indicates that Arf6 contributes to the β-arrestin-independent pathway. Collectively, our results clarify the contributions of GRKs and β-arrestins in FPR1 internalization, indicate that internalization involves multiple compensatory routes, and uncover previously unidentified regulators of FPR1 biogenesis and trafficking, providing new mechanistic insight into these processes.

## Introduction

Neutrophils serve as a primary defense against bacterial and fungal pathogens, but they also exacerbate inflammation and slow wound healing (Peiseler & Kubes, 2019). Indeed, excessive neutrophil activity is implicated in the pathogenesis of a broad range of inflammatory, autoimmune, and cardiovascular diseases (Herrero-Cervera et al., 2022). To mediate both their protective and damaging effects, neutrophils migrate from blood vessels through tissue to accumulate at sites of infection, injury, or inflammation. Their recruitment is controlled by a family of chemoattractant receptors on the cell surface that respond to a diverse set of pathogen- and host-derived molecules. After recruitment, these same receptors can also trigger other cytotoxic and inflammatory activities. Most chemoattractant receptors belong to a specialized subfamily of G protein-coupled receptors (GPCRs) that signal through Gαi G-proteins to direct cell migration (Metzemaekers et al., 2020).

The cell surface levels of chemoattractant receptors are actively regulated, which serves as a major mechanism to control the sensitivity and overall activity of neutrophils (Dahlgren et al., 2020). Stimulation of neutrophils through GPCRs and other receptors can cause mobilization of more chemoattractant receptors to the cell surface as part of a “priming” response that increases sensitivity (Dahlgren et al., 2023). Conversely, ligand binding by chemoattractant receptors triggers receptor internalization, which reduces signaling outputs and responsiveness to future stimuli, thereby helping to limit inflammatory responses (Dorward et al., 2015; Metzemaekers et al., 2020). Despite the importance of these processes, there are large gaps in our understanding of chemoattractant receptor internalization and cell surface expression.

The N-formyl peptide receptor FPR1 is a central regulator of neutrophil activity that serves as a primary model for chemotaxis signaling (He & Ye, 2017; Metzemaekers et al., 2020). FPR1 has been explored as a therapeutic target to limit neutrophil-dependent damage in inflammation (Honda et al., 2017; Lind et al., 2021) and as a candidate receptor for targeted drug delivery through its rapid internalization upon stimulation (Wang et al., 2019). While internalization of FPR1 could be therapeutically relevant, it does not seem to follow the best characterized pathways for endocytosis of GPCRs.

In the classic model of GPCR activation, ligand-dependent conformational changes in the receptor trigger GTP binding by Gα subunits, causing the dissociation of Gα-GTP and Gβγ and initiating a complex array of downstream signaling events (Pierce et al., 2002) (Figure 1A). Ligand-bound GPCRs get phosphorylated by GPCR kinases (GRKs), and phosphorylated receptors then interact with β-arrestins, which direct them to endocytic routes such as the canonical clathrin-mediated endocytosis pathway. Each of these three events, phosphorylation, β-arrestin binding, and internalization, can have critical effects for limiting inflammatory neutrophil functions through timely termination of G-protein signaling and desensitization of the receptors. The mechanisms that control FPR1 internalization are poorly understood, in part because it has been difficult to identify clear roles for components of known endocytosis pathways, including whether GRKs and β-arrestins are required.

**Figure 1.**
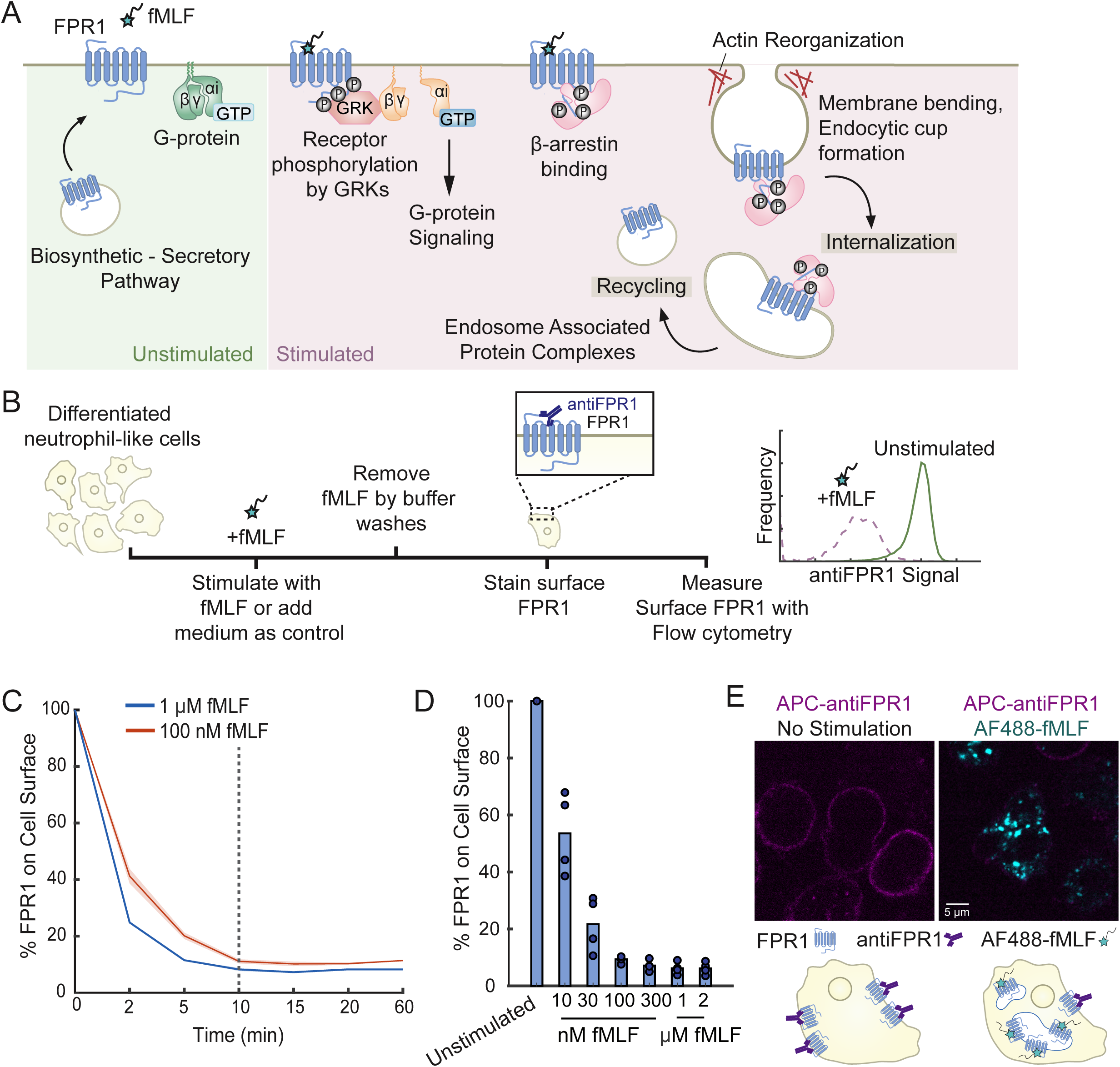
Characterization of FPR1 endocytosis. (A) Schematic summarizing FPR1 trafficking and key regulatory steps before and after fMLF stimulation. (B) Schematic of the flow cytometry- based FPR1 internalization assay. Differentiated neutrophil-like HL-60 cells are stimulated with fMLF to induce receptor internalization at 37°C. Remaining surface FPR1 is stained with an anti- FPR1 antibody on ice. The level of internalization is calculated relative to the FPR1 measurement of unstimulated cells. (C) Differentiated HL-60 cells were treated with 100 nM or 1 μM fMLF at different time-points (n=2, shaded region indicates mean +/- standard error). (D) Differentiated HL-60 cells were treated with different fMLF concentrations for 10 mins (n=4). (E) dHL-60 cells were either stimulated with AF488-fMLF or not stimulated for 5 min, fixed, and stained with an anti-FPR1 antibody (scale bar, 5μm).

There is evidence that GRKs regulate FPR1, but their role in controlling FPR1 endocytosis has remained unclear. First, fMLF stimulation of neutrophils from human blood leads to phosphorylation of FPR1 on seven of the eleven serine and threonine residues in its C-terminal tail (Maaty et al., 2013). Biochemical analysis implicated GRK2 activity in this process, as purified GRK2 and to a lesser degree GRK3 could phosphorylate the C-terminal tail of FPR1, while GRK5 and GRK6 had negligible activities (Prossnitz et al., 1995). GRK2 also gets recruited to FPR1 at the plasma membrane in stimulated cells (X. Liu et al., 2012), highlighting this kinase as a likely main regulator of FPR1 phosphorylation. However, GRK2 knockout neutrophil-like cells can still internalize FPR1 efficiently, which casts doubt on this role (B. C. Subramanian et al., 2018). Receptor phosphorylation is likely to be critical, as multiple publications from different groups have shown that a non-phosphorylatable FPR1 mutant is defective in internalization (Potter et al., 2006; Gilbert et al., 2001; X. Liu et al., 2012; B. C. Subramanian et al., 2018). Together, these results raise the question of whether GRKs other than GRK2 are involved. As a potential explanation for the biochemical results, the activities of GRKs can be primed. For instance, GRK2/3 phosphorylation of the chemokine receptor CXCR2 is necessary for binding of the Commander complex proteins COMMD3/8, which recruits GRK6 to the receptor (Nakai et al., 2019). GRK6 further phosphorylates this receptor to ensure β-arrestin2 binding (Nakai et al., 2019). Therefore, contribution of GRKs and receptor phosphorylation to FPR1 endocytosis is still an open question.

The role of β-arrestins in FPR1 internalization has also been unclear. Exogenously expressed FPR1 can be internalized in Mouse Embryonic Fibroblasts (MEFs) lacking β-arrestin1 and 2, but internalized receptors accumulate in endosomes (Vines et al., 2003). These results suggest that β-arrestins might be involved in FPR1 recycling but not in its internalization. However, both β- arrestin1 and 2 are recruited to the plasma membrane in neutrophil-like cells in response to fMLF stimulation (Wang & Ye, 2022). If we consider the discrepancies between cell types, mouse and human FPRs (Winther et al., 2018), and the importance of phosphorylation patterns in β-arrestin binding and receptor control (Mayer et al., 2019), it is unclear how β-arrestins regulate FPR1 surface expression in neutrophils.

FPR1 internalization does not require the main endocytic adaptor proteins clathrin or flotillin (Gilbert et al., 2001; B. C. Subramanian et al., 2018). Caveolins are not expressed in primary human neutrophils or in differentiated neutrophil model cell lines (Rincón et al., 2018). Although these negative results strongly suggest that FPR1 uses multiple or uncommon endocytic pathways, there is a need for systematic and unbiased approaches to identify the unknown regulators of FPR1 endocytosis. Moreover, the regulation of FPR1 forward trafficking has also been unclear. FPR1 is synthesized during late stages of neutrophil differentiation, but its surface secretion is controlled. A sizable fraction of these receptors are stored in secretory vesicles and granules that can be mobilized to the plasma membrane in response to inflammatory stimuli (Dorward et al., 2015; Sengeløv et al., 1994).

Pooled CRISPR screening is a powerful technology to identify genes and associated pathways that regulate different cellular processes (Bock et al., 2022). CRISPR-based screens outperform previous technologies based on RNA interferences, with higher sensitivity and lower rates of false positive hits from off-target activity (Evers et al., 2016; Morgens et al., 2017). CRISPR systems using the Cas9 endonuclease or the catalytically inactive Cas9 (dCas9) with single-guide RNA (sgRNA) libraries targeting every protein-coding gene in the human genome have been used as discovery tools for many biological questions including regulation of GPCR trafficking (Novy et al., 2024), phagocytosis (Haney et al., 2018), and neutrophil migration and differentiation (Belliveau et al., 2023).

Here, we investigated the roles of both classic and unknown endocytosis regulators. Although their roles had been unclear, we found that multiple GRKs and both β-arrestin1 and 2 are important for FPR1 internalization. Moreover, lack of β-arrestins cause partial defects in FPR1 internalization, strongly suggesting that multiple internalization routes are used by this receptor. We performed two parallel genome-wide CRISPR/Cas9 screens for the regulators of FPR1 surface expression and internalization. Taking advantage of the data collected from multiple guides per gene and the tunable nature of our cell selection strategy, we were able to capture hits with mild and more severe phenotypes. We integrated data from the two screens to identify hits that are specific to basal FPR1 surface expression or its regulation post-stimulation. We found that disruption of multiple multi-protein complexes with known roles in receptor recycling cause defects in different steps of FPR1 trafficking. We validated two internalization-specific hits whose roles were previously unrecognized in FPR1 endocytosis: the formin diaphanous 1 (mDia1; also known as DIAPH1) and the small GTPase Arf6. We showed that chemical inhibition of their activities causes significant defects in FPR1 internalization in both neutrophil-like cells and primary human neutrophils. Our work not only identifies new regulators of FPR1 trafficking but also provides a useful dataset that has the potential to advise future studies on GPCR trafficking, as well as efforts to develop therapies based on controlling receptor activity or surface availability.

## Results

### Ligand-induced FPR1 endocytosis is fast and efficient

FPR1 endocytosis is typically investigated by the quantification of the internalized fluorescent formyl peptides or exogenously expressed fluorescently tagged FPR1 using flow cytometry and confocal microscopy (B. C. Subramanian et al., 2018; Vines et al., 2003). These techniques have been valuable in studying endocytic mechanisms of various receptors, but the throughput of microscopy is limited, and in each case, it can be difficult to discriminate precisely between receptors that have been internalized and those still on the surface (leading to higher background). To address these challenges, we optimized a flow-cytometry based FPR1 internalization assay in which we measure FPR1 surface expression using immunostaining, comparing unstimulated cells with cells stimulated with fMLF (Figure 1B-D). The decrease in mean fluorescence signal in stimulated cells serves as a measure of FPR1 internalization and recycling. To validate our approach, we used confocal microscopy to monitor internalization of a fluorescent ligand (Alexa Fluor 488 conjugated fMLF or AF488-fMLF) and immunostaining of surface FPR1 in individual cells. We observed decreases in surface FPR1 staining that agrees with the corresponding signal of internalized chemoattractant (Figure 1E). Importantly, our assay accounts for the variations in basal FPR1 surface expression that can be caused by the differentiation efficiency, small chemical inhibitor or siRNA treatments, and gene editing.

Since it is not feasible to make gene-specific knockdowns in primary human neutrophils, we used the neutrophil-like cell line HL-60 to study FPR1 endocytosis. Where feasible, we confirmed our findings in primary human neutrophils. As an initial characterization, we found that FPR1 internalization depends quantitatively on both time and fMLF concentration (Figure 1C and D). Following a 2-minute stimulation of 100 nM fMLF, more than 50% of surface FPR1 is internalized (Figure 1C), and by 10 minutes only around 20% of the initial FPR1 remains on the surface (Figure 1C and D). These results indicate that ligand-induced FPR1 internalization is fast and efficient, consistent with previous findings (Dorward et al., 2015). Even so, we observed a maximum internalization capacity for FPR1 of ∼80-90%. Longer incubations or higher fMLF concentrations did not lower the remaining surface FPR1 levels in differentiated HL-60 cells. This result suggests that robust recycling of internalized receptors may partially offset endocytosis. Since the maximum internalization is achieved by 10 minutes of fMLF stimulation, we choose this incubation time for future experiments to investigate the components of the FPR1 endocytic mechanism. Additionally, since FPR1 surface expression can be heterogeneous in neutrophil-like cell lines (Rincón et al., 2018), we use differentiated HL-60 cells that exogenously express a 3XHA-FPR1 construct to achieve a uniform FPR1 expression for further investigation. These cells can internalize surface FPR1 at similar levels to differentiated wild-type HL-60 cells (Supplementary Figure 1A and B), indicating that the 3XHA tag does not interfere with receptor trafficking. For simplicity, we refer to differentiated cells expressing the exogenous receptor simply as dHL-60s.

### Multiple GRKs contribute to FPR1 endocytosis

The role of GRKs in controlling FPR1 endocytosis has been unclear. GRK2 can phosphorylate FPR1 (Prossnitz et al., 1995), but it is not required (B. C. Subramanian et al., 2018). However, FPR1 with a mutated non-phosphorylatable C-terminal tail causes defects in internalization (Gilbert et al., 2001; X. Liu et al., 2012; Potter et al., 2006; B. C. Subramanian et al., 2018). Thus, GRKs other than GRK2 may be involved in FPR1 phosphorylation and internalization. Four GRKs are expressed in neutrophils: GRK2, GRK3, GRK5, and the lymphoid-enriched GRK6 (Tiedemann et al., 2010). To resolve the roles of GRKs in FPR1 internalization, we analyzed the effects of losing function of these four proteins, both individually and in combinations.

Since GRK3 is closely related to GRK2, sharing similar regulation by G-proteins (Willets et al., 2003), we used the small molecule inhibitor Compound 101 (GRK2/3 inhibitor) to block the function of these two kinases simultaneously. Although GRK2/3 inhibition led to a significant defect in FPR1 internalization, most of the surface FPR1 (>50% at 100 nM fMLF and >75% at 1 μM fMLF) can still be internalized in these cells, even at high inhibitor concentrations (Figure 2A). This result indicates that GRK2 and GRK3 contribute to FPR1 endocytosis, but also that additional kinases are likely involved.

**Figure 2.**
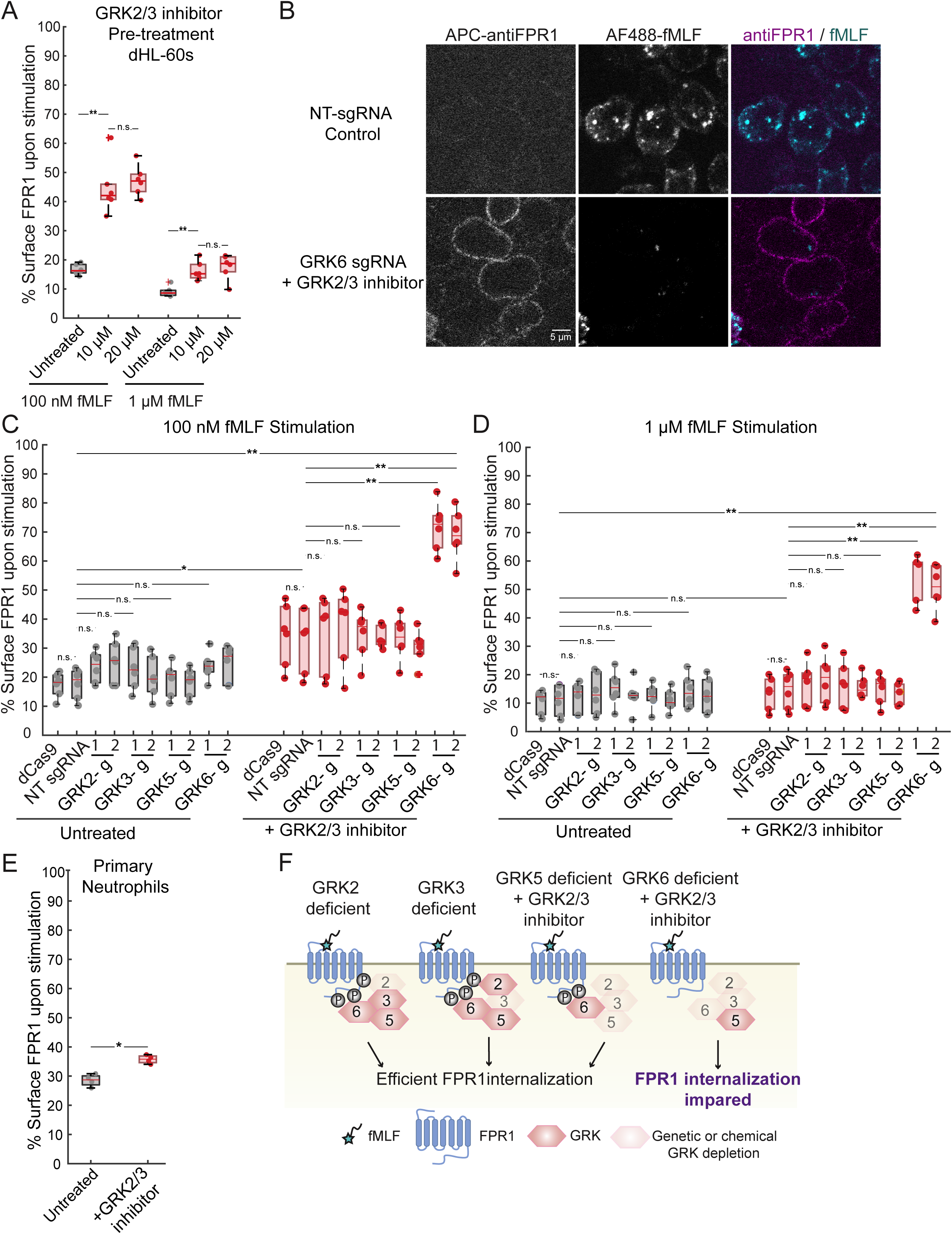
GRK2, GRK3 and GRK6 take roles in FPR1 endocytosis. (A) Cells were untreated or pre-treated with the GRK2/3 inhibitor Compound 101 (10 or 20 μM) for 30 min and then stimulated with either 100 nM or 1 μM fMLF for 10 min (n=6). (B) dCas9 expressing dHL-60s (Control) and Compound 101 pretreated dHL-60 cells expressing dCas9 and GRK6 sgRNA#1 were stimulated with AF488-fMLF for 5 min, fixed, and stained with an anti-FPR1 antibody (scale bar, 5μm). Control and CRISPRi knockdown cells were untreated or pre-treated with Compound 101 for 30 min and then stimulated with either (C) 100 nM or (D) 1 μM fMLF for 10 min (n=6). ‘g’ stands for sgRNA, and 1 and 2 are unique guide sequences. (E) Primary human neutrophils were untreated or pre-treated with 20 μM Compound 101 for 30 min and then stimulated with 100 nM fMLF for 10 min (n=4). Throughout this study, we use Mann-Whitney U test for pairwise comparison of the mean fluorescence signal (bar height) of the two samples indicated in figures (n.s. p>0.05, *p<0.05, **p<0.01, ***p<0.001), and dots represent different biological replicates. (F) Schematic summary of how defects in different GRK activities affect FPR1 internalization.

To test the roles of each GRK systematically, and to analyze their combined roles with GRK2 and GRK3, we generated cell lines with stable CRISPRi knockdowns of GRK2, GRK3, GRK5, and GRK6. Each knockdown cell line had a >70% reduction in the target gene expression, except one of the GRK2 knockdowns (sgRNA#1) which had >60% reduction in GRK2 mRNA levels (Supplementary Figure 2). We measured FPR1 internalization efficiency for each cell line, with and without the GRK2/3 inhibitor, and confirmed our key results using confocal microscopy (Figure 2A-D). We detected no significant defects in FPR1 internalization in any of the single GRK knockdown cell lines (Figure 2C and D). However, treatment of the knockdown cell lines with the GRK2/3 inhibitor revealed significant defects. Inhibitor treatment of cells with GRK6 knockdown, but not GRK5, led to a dramatic decrease in FPR1 internalization upon stimulation (less than 30% of surface FPR1 internalized with 100 nM fMLF), suggesting that the combined activities of GRK2, GRK3, and GRK6 are key for FPR1 internalization (Figure 2C and F). At a higher fMLF concentration, defects caused by the GRK2/3 inhibitor alone were less evident (Figure 2A and D), but the cells with reduced GRK2, GRK3, and GRK6 activities still showed a significant defect in FPR1 internalization (Figure 2D). Using confocal microscopy, we confirmed that AF488-fMLF internalization was minimal in GRK2/3/6 deficient cells compared to the control cells, and GRK2/3/6 deficient cells retained FPR1 on their surface after stimulation (Figure 2B).

We tested the effect of the GRK2/3 inhibitor in primary human neutrophils and detected a significant but partial defect in FPR1 internalization (Figure 2E), consistent with our findings in dHL-60 cells.

Collectively, these results demonstrate the involvement of multiple GRKs in FPR1 endocytosis, including GRK6 (Figure 2F), and strongly support a requirement for ligand-induced receptor phosphorylation for FPR1 internalization.

### FPR1 uses both β-arrestin-dependent and β-arrestin-independent endocytic pathways

Although β-arrestins control the endocytosis of many GPCRs, FPR1 can internalize in a β- arrestin-independent manner, at least in heterologous cell types. A mutant FPR1 defective in binding β-arrestins can still internalize in HEK-293 cells (Bennett et al., 2000), and FPR1 can internalize in mouse embryonic fibroblasts lacking β-arrestins (Vines et al., 2003). Even so, β- arrestins are recruited to FPR1 upon fMLF stimulation (Wang & Ye, 2022), and their function in human neutrophils needs further investigation.

To test if FPR1 internalization requires β-arrestin activity in a neutrophil model, we generated single (β-arrestin1 or 2) and double (β-arrestin1 and 2) CRISPR knockout cell lines, initially as uncloned cell pools. Using the flow cytometry-based internalization assay, we found that FPR1 internalization efficiency is significantly decreased in single β-arrestin knockouts: ∼10-15% more FPR1 was retained on the cell surface post-stimulation in each of the single knockouts (Figure 3A and B). These defects were synergistic, causing a more severe phenotype in β-arrestin double knockouts (DKOs) (∼50-60% reduction in FPR1 internalization capacity) (Figure 3A and B). However, consistent with previous studies, FPR1 can still be internalized in the absence of β-arrestins. β-arrestin DKOs can also internalize AF488-fMLF efficiently, but we observed dimmer puncta in these cells (Figure 3C). These findings suggest that FPR1 uses both β-arrestin-dependent and -independent endocytic pathways (Figure 3D).

**Figure 3.**
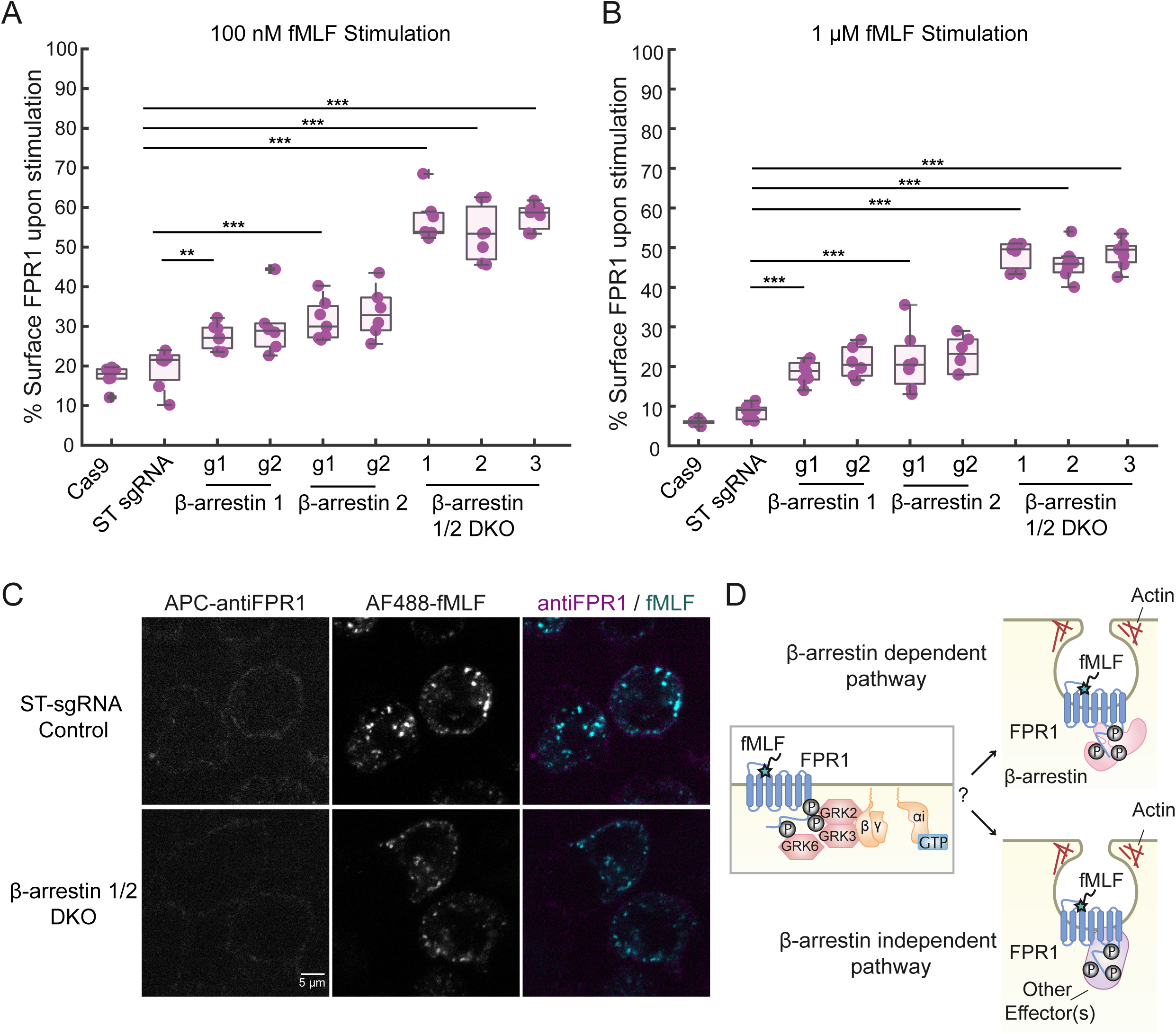
β-arrestin1 and 2 take roles in FPR1 endocytosis. Cells were stimulated with either (A) 100 nM or (B) 1 μM fMLF for 10 mins. Dots represent different biological replicates (n=6 for Cas9, β-arrestin1 g2, and β-arrestin2 g2, and n=7 for safe-targeting (ST) sgRNA, β-arrestin1 g1, and β-arrestin 2 g1). ‘g’ stands for sgRNA, and 1 and 2 are different guide sequences. (C) ST sgRNA expressing dHL-60s (Control) and β-arrestin double knockout cells were stimulated with AF488-fMLF for 5 min, fixed, and stained with an anti-FPR1 antibody (scale bar, 5μm). (D) Schematic representation of potential β-arrestin dependent and independent endocytic routes used by FPR1.

To rule out the possibility that heterogeneity in the pooled CRISPR knockout cell lines was responsible for the incomplete blockage of FPR1 internalization in β-arrestin1/2 DKOs, we isolated multiple clones from the pooled DKO cell population. The cloned cell lines internalize FPR1 with similar defects to those seen for the pooled cell lines (Supplementary Figure 3). Moreover, we used amplicon sequencing to determine the mutations in the pooled DKO cell line and three of the clones. For the pooled cell line, more than 89% of reads for β-arrestin1 indicated mutations and more than 97% of the reads for β-arrestin2, with high frequencies of deletion and frameshift mutations. These results confirmed the high efficiency of the knockouts in pooled cell lines (Supplementary Figure 4).

### Design and analysis of two parallel genome-wide screens for regulators of FPR1 endocytosis and surface expression

To provide insight into the unknown modulators of FPR1 surface expression before and after fMLF stimulation, we performed two parallel CRISPR deletion screens (Figure 4A). We constructed an HL-60 cell line that stably expresses a 3XHA-FPR1 construct and the Cas9 nuclease, and we verified that Cas9 works efficiently in these cells using a “self-targeting” GFP construct (Supplementary Figure 1C). We then infected these cells with a genome-wide CRISPR knockout library that consists of 10 sgRNAs per gene, as well as 5644 non-targeting and 6750 safe-targeting sgRNAs (Morgens et al., 2017). To maintain library representation, we aimed to have at least 1,000 cells per guide during infection and subsequent passaging steps. To complete the screens, we differentiated ∼300 million cells (1000X coverage) for both unstimulated and fMLF-stimulated conditions. After fMLF or mock stimulation, we incubated cells for 10 minutes to allow endocytosis and stained for surface FPR1 (Figure 4A). Using Fluorescence-activated cell sorting (FACS), we collected the top and bottom 30% of the anti-FPR1 stained cells for each sample. We then determined the abundance of the sgRNAs expressed in these populations using next-generation sequencing.

**Figure 4.**
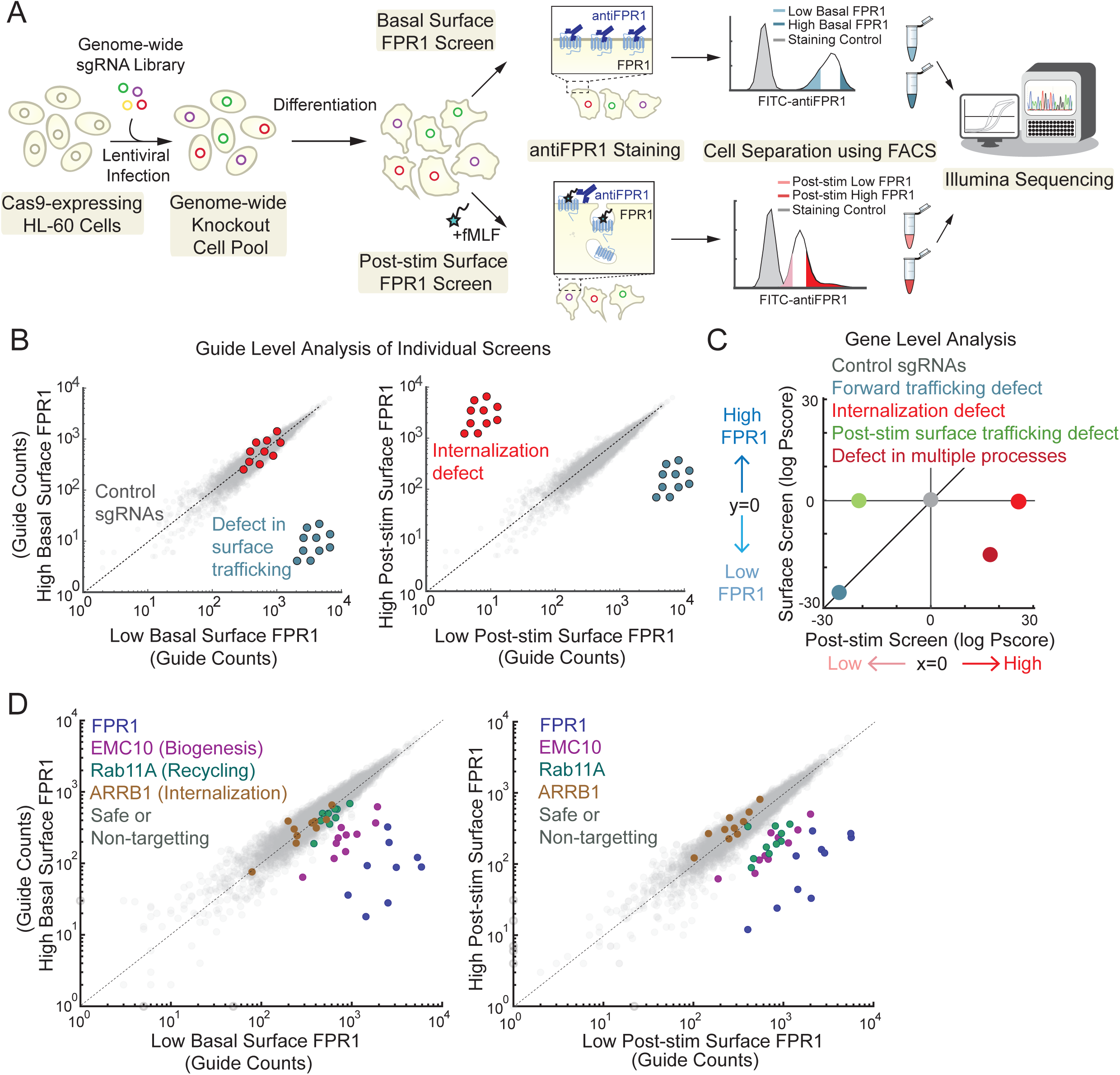
Overview of genome-wide CRISPR/Cas9 screen design and expected hits. (A) Schematic representation of the two parallel screen designs for the regulators of FPR1 basal and post-stimulation surface expression. Example scatter plots showing where hits from the two screens are expected to be found (B) at the guide level (comparing guide counts from different samples) and (C) at the gene level (comparing log P-score for each gene in two screens). (D) Scatter plots showing unique guide counts of FPR1, three expected hits, EMC10 (biogenesis), RAB11A (recycling), and ARRB1 (internalization), and distribution of control guides.

In combination, we reasoned that the two bidirectional screens would allow identification and discrimination of genes whose loss-of-function affects cell surface FPR1 expression in different ways, including gene expression, secretion, internalization, and recycling. For example, guides targeting genes needed for FPR1 gene expression, biogenesis, or trafficking to the cell surface should be enriched in the low bins in both screens, whereas guides targeting genes required for internalization should be enriched in the high bin specifically in the post-stimulation screen (Figure 4B). Comparing gene-level scores from the two screens should distinguish these phenotypes from defects in recycling or more complex phenotypes in which multiple processes are affected (Figure 4C).

In contrast to most genome-wide screens, determining significance for the range of phenotypes observable in our screens required joint analysis of all four samples. To do this while accounting for read-level statistical noise, we developed a log-likelihood scoring system based on a negative binomial model. We combine guide-level scores to obtain gene-level scores, and we compute p-values using a permutation-based approach that takes advantage of the large number of control guides, similar to the approach used in the casTLE algorithm (Morgens et al., 2017). Finally, we estimate effect sizes using a Bayesian approach (see Methods for more details). We thus computed statistical scores, effect size estimates, and p-values for every gene for basal FPR1 surface expression, post-stimulus FPR1 surface expression, and for relative differential FPR1 surface expression between the two conditions (Supplementary Table 1). To compute formal lists of “hits” and for downstream analyses, we used a logP score which is the log10 p-value multiplied by the sign of the effect.

As an initial validation of our screens, we examined the guide-level data for genes with known roles in biogenesis, recycling, and internalization, and we found that they were enriched in different bins, consistent with their roles. Guides targeting FPR1 were enriched in the bins for low surface expression in both screens (Figure 4D). We observed similar enrichments for EMC10, which is a member of the ER membrane protein complex (EMC) that is important for proper protein folding and membrane localization of transmembrane proteins (Guna et al., 2018; Shurtleff et al., 2018) (Figure 4D). More generally, our screens identified eight members of this complex as regulators of FPR1 biogenesis and surface localization, independent of stimulation. The small GTPase Rab11 is a key regulator of vesicular trafficking with known functions in stimulation-dependent FPR1 recycling (B. C. Subramanian et al., 2018; Vines et al., 2003). The guides targeting Rab11A (as well as those targeting its paralog Rab11B) were enriched in the low surface expression bin specifically after stimulation, consistent with a role in recycling (Fig. 4D). Finally, the guides targeting β-arrestin1 (ARRB1) were modestly biased towards the high bin in the post-stimulation samples. While this was the direction expected for an internalization hit, the modest size of the effect may be due to its overlapping role with β-arrestin 2 and the lack of a strict requirement for β-arrestins in FPR1 internalization.

### Parallel genome-wide CRISPR/Cas9 screens reveal diverse regulators of FPR1 biogenesis and trafficking

In total, we identified 2507 hits across the different phenotypes: 849 genes positively (knockout causes low basal surface FPR1) and 399 genes negatively (knockout leads to high basal FPR1) regulate basal FPR1 surface expression, while 515 genes negatively and 744 genes positively affect FPR1 surface expression post-stimulation (Figure 5A and B, Supplementary Tables 2-5). Through integrated analysis of the two screens, we identified 85 negative (knockout enhances FPR1 internalization or downregulates recycling, resulting in low surface FPR1) and 277 positive regulators of FPR1 internalization (knockout decreases FPR1 internalization or enhances recycling, resulting in high surface FPR1) (Figure 5A and B, Supplementary Tables 6 and 7).

**Figure 5.**
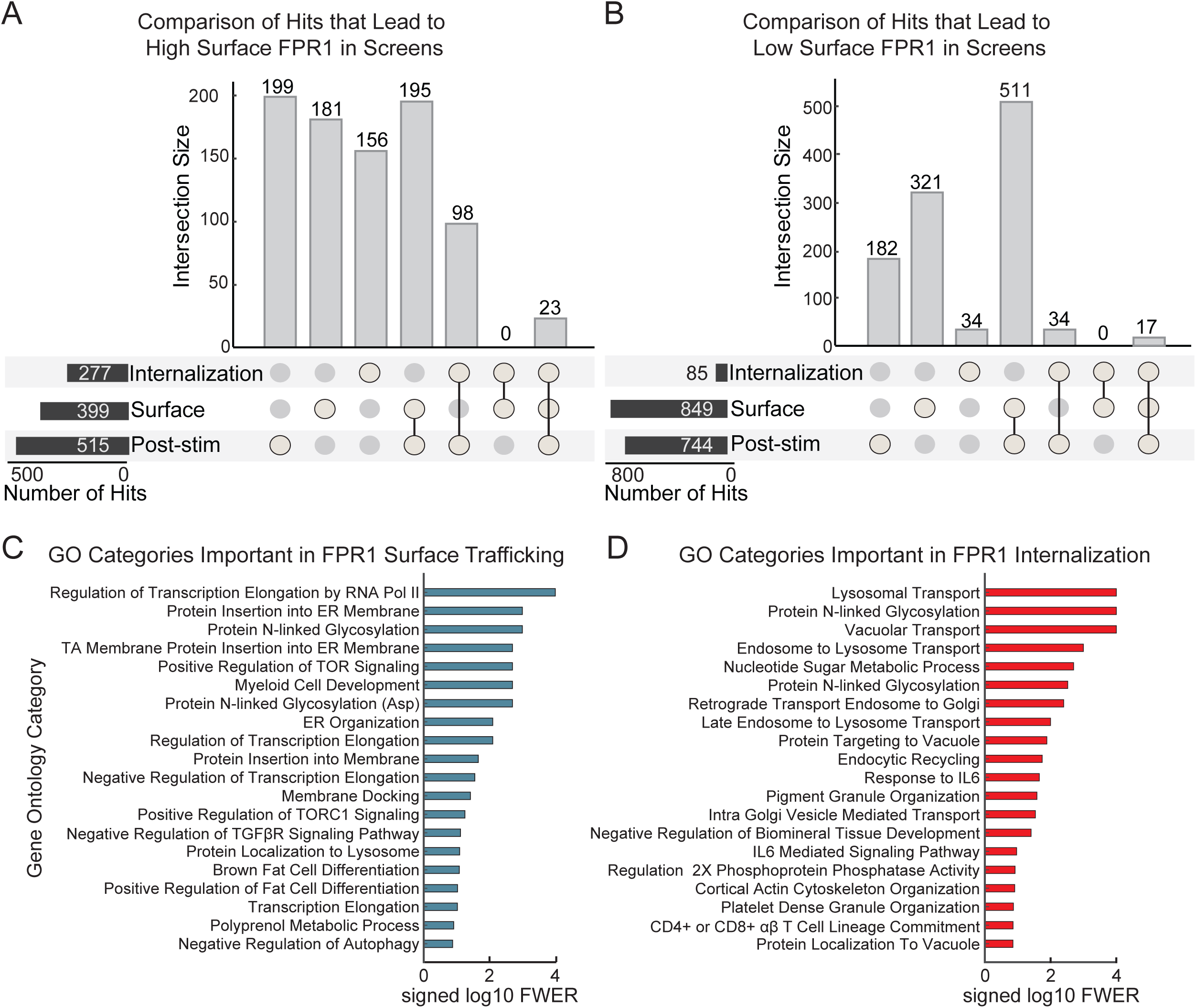
Summary of the number of hits across different phenotypes obtained from the CRISPR screens and GSEA. UpSet plots (Conway et al., 2017) showing distribution of the number of hits that negatively (A) or positively (B) regulate FPR1 surface expression in the basal state or after fMLF stimulation (hits were determined using a one-tailed pFDR< 0.05 threshold). Vertical bars represent the intersection between the indicated sets, or the unique members of a single set. Horizontal (black) bars represent the total number of hits for each screen or integrated analysis. Gene set enrichment analysis (Reimand et al., 2019) summarizing the top 20 significant gene ontology categories for the hits from the FPR1 basal surface expression screen (C) and the integrated analysis for FPR1 internalization hits (D).

To get an overview of the genes identified, we performed gene set enrichment analysis (GSEA) (A. Subramanian et al., 2005), focusing on the basal surface FPR1 screen and the integrated analysis of internalization and recycling. For low basal surface FPR1 expression, the most significantly enriched gene sets were involved in transcription regulation, ER processing, membrane insertion, N-linked glycosylation, TOR signaling, and cell differentiation (Figure 5C). The hits for FPR1 internalization obtained from the integrative analysis were enriched for gene sets including regulators of organelle transport, endocytic recycling, and cortical actin (Figure 5D).

To get a more detailed picture of the pathways involved, we manually curated hits with roles in signaling and trafficking pathways likely to directly regulate FPR1 surface localization (Figure 6A). Our screens identified components of GPCR signaling, protein folding and sorting machinery, the cytoskeleton, and intracellular trafficking.

**Figure 6.**
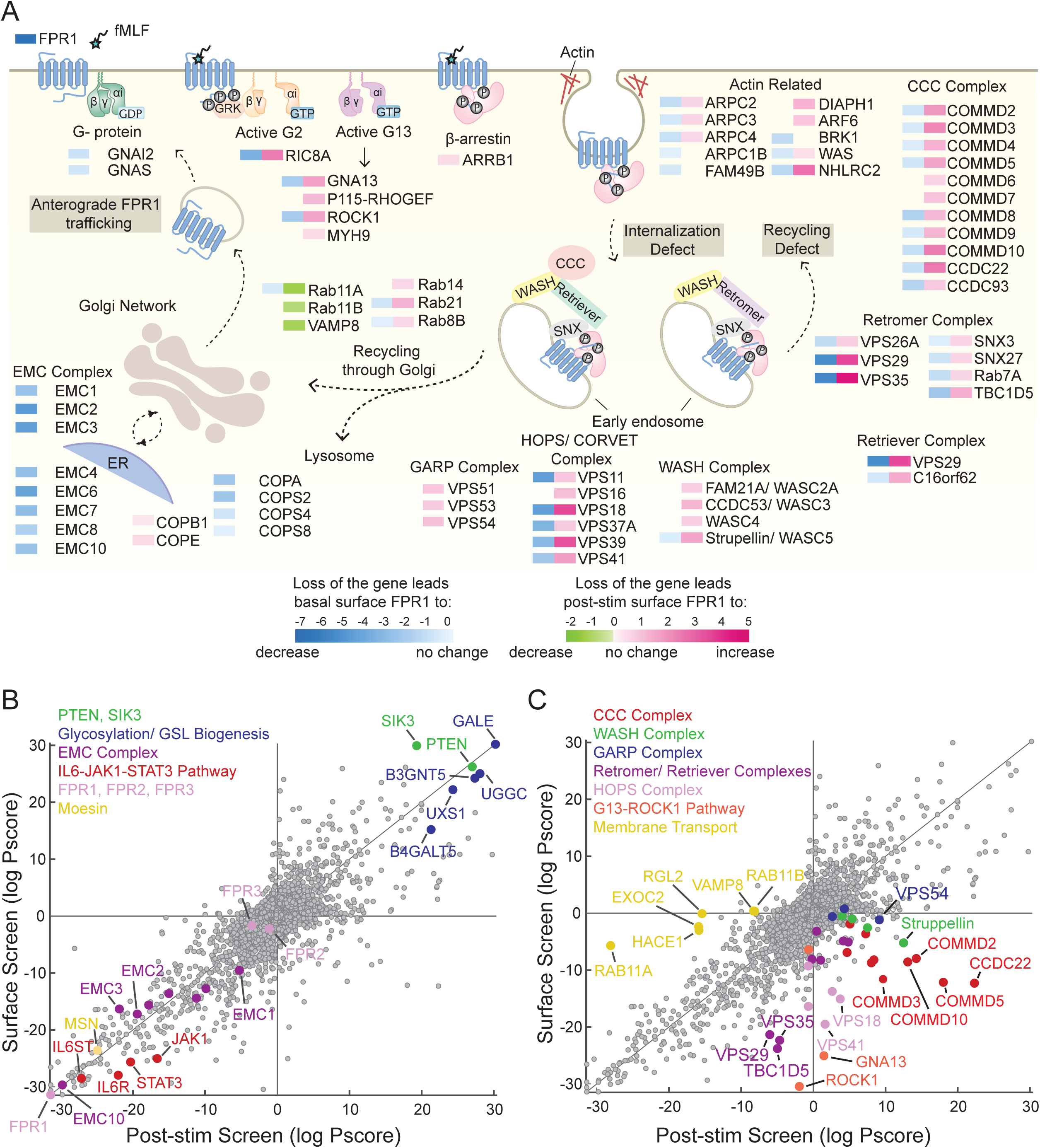
Summary of the hits from the parallel whole-genome CRISPR screens for the regulators of FPR1 surface expression, endocytosis, and recycling. (A) Schematic representation of a selection of hits found in the parallel whole-genome CRISPR screens. Genes shown are significant hits in at least one screen, and they were organized based on prior literature. The effect scores for each gene are represented as colored rectangles. The scores are only shown if the gene was a hit in the particular screen (pFDR< 0.05). Negative scores indicate a decrease in surface FPR1, while positive scores indicate an increase. For hits regulating FPR1 surface expression post-stimulation (shown in pink and green), the effect scores obtained from the integrated analysis of the two screens were presented. (B) Top hits from the surface FPR1 expression screen. FPR2 and FPR3 were shown for comparison and are not identified as hits in either of the screens. (C) Significant hits from the integrated analysis of FPR1 internalization, recycling, or exocytosis. Scatter plots in B and C represent the same dataset (log Pscore, with the basal surface expression score on the y-axis and post-stimulation score on the x-axis) with different genes highlighted.

### Genome-wide CRISPR/Cas9 screens reveal regulators of basal FPR1 surface expression

The genes affecting basal surface FPR1 expression include general regulators of cell cycle, transcription, translation, protein biogenesis and trafficking, and neutrophil differentiation.

Our screen revealed multiple genes that are critical for efficient trafficking of FPR1 to the cell surface (Supplementary Table 2). In addition to EMC components, COPI coat proteins (COPA in ER- Golgi retrograde transport), components of the TRAPP (TRAPPC2L, TRAPPC13) and Exocyst complexes that function in vesicle tethering (EXOC2, EXOC3L2), and a member of ezrin/radixin/moesin (ERM) protein family, moesin (MSN), that links F-actin to plasma membrane appear to be important for FPR1 biogenesis and surface secretion (Figure 6A, B and Supplementary Table 2). Arp2/3 complex components and an Arp2/3 activator, the Wiskott– Aldrich syndrome protein (WASP), were also among the hits, highlighting the importance of an active cytoskeletal machinery for FPR1 trafficking (Figure 6A and 7A).

**Figure 7.**
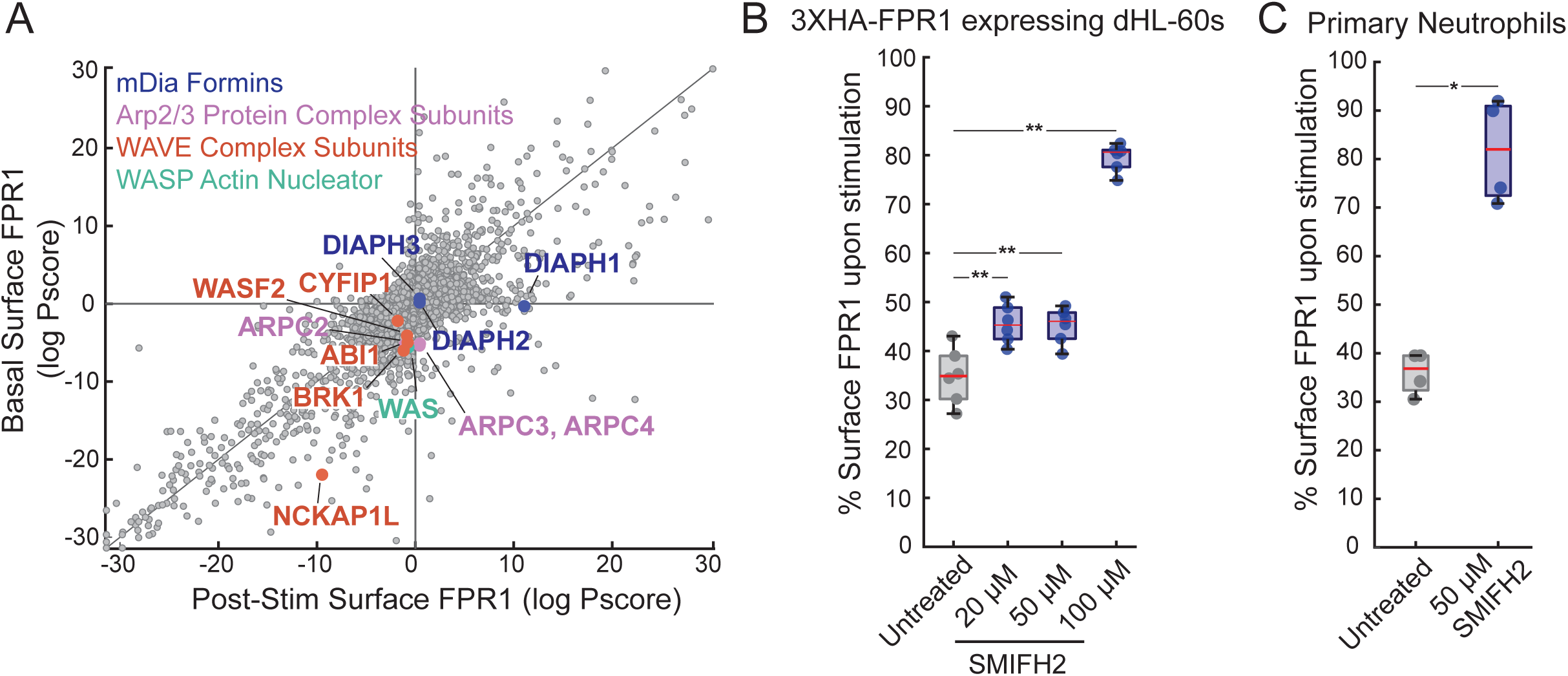
Fomin activity is important for efficient FPR1 internalization. (A) Plot showing the effect of the knockout of major actin regulators on FPR1 surface levels in the basal (y-axis) and post-stimulation screens (x-axis). Effect of SMIFH2 on FPR1 internalization (B) in dHL-60s (20 μM, 50 μM, and 100 μM) and (C) in primary neutrophils (50 μM). Cells are untreated or pre-treated with SMIFH2 for 20 mins. (n=6 for dHL-60s, n=4 for primary neutrophils).

FPR1 expression is also correlated with neutrophil differentiation, as it appears on the cell surface at later stages of differentiation (Rincón et al., 2018). Consistent with this, we found that targeting mediators of neutrophil differentiation caused decreased basal surface FPR1 levels. Some of the most significant hits were TOR complex components and regulators of TOR activity, including MLST8, RRAGA, RICTOR, LAMTOR2/4, CSNK1A1, and FLCN (Supplementary Figure 5). These findings agreed with a recent genome-wide CRISPRi screen for genes controlling differentiation of HL-60 cells, which identified both mTORC1 and mTORC2 signaling as central for differentiation (Belliveau et al., 2023). Similarly, transcription factors involved in differentiation to mature granulocytes (CAAT/enhancer binding proteins, CEBPA/B/D/E, and PU.1 also known as SPI1) and genes that function in the maturation of SUMO proteins (SENP1/2/3 and SUMO2) were shared hits for our basal surface FPR1 screen (Supplementary Figure 5) and the earlier differentiation screen (Belliveau et al., 2023). Our screen also revealed other potential differentiation regulators that were not identified in the differentiation screen, including IL6R, IL6ST, STAT3, and JAK1 (Figure 6B). Although IL6 was reported to be important for B-cell (Joshizaki et al, 1984) and dendritic cell maturation (Park et al., 2004), its roles in FPR1 trafficking and neutrophil differentiation are not clear. Our screen suggests an important role for the IL6/JAK1/STAT3 pathway in FPR1 surface expression, but it remains to be determined if this signaling pathway is important for late stages of neutrophil differentiation or FPR1 trafficking in particular.

On the other hand, we identified genes that limit FPR1 surface expression (Supplementary Table 3), including the two previously annotated negative regulators of neutrophil differentiation (Gene Ontology Annotation, GO: 0045659), TRIB1 and INPP5D (also known as SHIP1). Loss of tumor suppressor genes PTEN and STK11 (also known as LBK1) and the STK11-regulating kinases (SIK1, SIK3) also elevates basal surface FPR1 levels (Figure 6B). Similarly, knockout of well-known genes involved in glycosphingolipid (GSL) biogenesis (UGCG, B4GLT5, B3GNT5) and N-linked glycosylation (GALE, UXS1) caused higher basal surface FPR1 levels (Figure 6B). As GSLs are crucial for forming GSL enriched microdomains (GEMs) that take diverse roles in immune cells (Zhang et al., 2019), it is possible that these domains exclude FPR1 or participate in the turnover of FPR1 at the membrane.

### Genome-wide CRISPR/Cas9 screens reveal regulators of ligand-induced FPR1 transport to plasma membrane

Genes whose knockout causes lower surface FPR1 after chemoattractant stimulation are potential regulators of ligand-induced FPR1 recycling or stimulation-dependent movement of FPR1 from intracellular vesicles to the plasma membrane (Supplementary Tables 4 and 6). We identified vesicle-associated membrane protein 8 (VAMP8), a SNARE protein involved in vesicle fusion with the plasma membrane, as a hit in this group (Figure 6A), suggesting a role for VAMP8 in mobilization of FPR1-containing granules in response to fMLF.

As mentioned earlier, Rab11 is known to regulate FPR1 recycling, and both Rab11 isoforms A and B were significant hits (Figure 6A and C). HACE1, a HECT-domain-containing ubiquitin ligase, which is known to promote recycling of other GPCRs by ubiquitinating Rab11 (Lachance et al., 2013) was also a hit in this group (Figure 6C). Rab11 can coordinate secretion through the exocyst complex, which mediates the delivery and tethering of secretory vesicles to specific sites on the plasma membrane (Bai et al., 2022). Vesicle tethering is also regulated through interactions of the exocyst with Ral GTPases (Shirakawa & Horiuchi, 2015). In our screens, loss of the exocyst component EXOC2 or the Ral GTPase regulator RGL2 caused significantly lower levels of surface FPR1 in stimulated cells (Figure 6C). In platelets, a defect in EXOC2-Ral GTPase interaction was shown to inhibit ligand-induced granule secretion (Kawato et al., 2008). These proteins may also be working together to facilitate fusion of FPR1-loaded vesicles with the plasma membrane in neutrophils.

### Integrated analysis of screens identifies multi-protein complexes and signaling components that regulate multiple aspects of FPR1 surface expression

Our integrated analysis of the two screens was designed to identify key factors for FPR1 endocytosis and recycling (Supplementary Table 7). The majority of our hits from this analysis had relatively increased surface FPR1 levels after stimulation, consistent with a role in endocytosis or limiting recycling. Top hits in this class include (1) the CCC (COMMD/CCDC22/CCDC93) Complex, which takes important roles in membrane protein recycling, (2) the WASH (Wiskott–Aldrich syndrome protein and SCAR homologue) Complex, which activates the Arp2/3 complex and induces F-actin branching, (3) the retromer/retriever complexes, which are important in cargo identification and its proper sorting, (4) the GARP (Golgi-associated retrograde protein) Complex, which facilitates endosome-Golgi retrograde transport (5) the HOPS (Homotypic Fusion and Protein Sorting) complex, which is important in endo- lysosomal organization, and involve (6) actin regulators and (7) signaling components (Figure 6A and C).

Nine genes that constitute the CCC complex were identified as regulators of FPR1 basal surface expression and endocytosis: COMMD proteins 2, 3, 5, 6, 7, 8, and 10 as well as CCDC22 and CCDC93. The CCC complex collaborates with other multiprotein trafficking complexes, including WASH and retriever, for cargo sorting (Laulumaa & Varjosalo, 2021). Consistent with the involvement of this broader pathway, we also identified components of the WASH (Strumpellin/KIAA0196/WASHC5, KIAA1033/WASHC4, CCDC53/WASHC3, and FAM21A/WASHC2A) and retriever (VPS29 and C16orf62) complexes (Figure 6A and C). Notably, mutation of most components of all three complexes caused decreased basal surface FPR1, in addition to the effect on endocytosis or recycling. Similarly, all three constituents of the retromer (VPS35, VPS26A, and VPS29) and the adaptor proteins important for retromer recruitment to endosomes (Rab7A and SNX3) were among hits for both screens. Although these complexes have similar roles in the cell, different integral membrane proteins may prefer to use the retromer or the retriever for recycling back to the cell surface (Gershlick & Lucas, 2017; D. Liu et al., 2022). Our screens point to a role for both retromer and retriever complexes in FPR1 sorting in neutrophils.

In addition to the trafficking complexes, we found multiple hits in the G13/RhoA/ROCK1 signaling pathway as potential regulators of endocytosis or recycling (Figure 6A and C). In response to fMLF stimulation, different heterotrimeric G proteins are activated including G12 and G13 (Xu et al., 2003). A well-studied effector of G12/13 is the GTPase RhoA, whose activity is linked to regulation of cortical actin reorganization at the trailing edge of polarized neutrophils (Hind et al., 2016; Xu et al., 2003). Moreover, G13-dependent signaling was shown to be important for the localization of a chemokine receptor CXCR4 to Rab11+ recycling endosomes (Kumar et al., 2011). This signaling pathway may have roles in locating FPR1 to the correct recycling endosomes.

### Formin activity regulates efficient FPR1 internalization

One of our primary goals for the screens was to gain new insights into the mechanisms controlling FPR1 internalization. Because many of the hits mentioned above are members of known endolysosomal complexes or are signaling components whose knockout can affect multiple cellular processes, we looked carefully for other genes that were hits specifically in the internalization screen. Among such genes, we selected the formin mDia1, which plays a critical role in actin filament formation, as an interesting hit.

Actin is involved in multiple endocytic pathways, but it is not required for all pathways, and the actin structures involved can vary. In some pathways, actin filaments are critical for the assembly of the endocytic machinery at the nascent vesicle buds, and actin can participate in vesicle scission in late stages of receptor internalization (Abouelezz & Almeida-Souza, 2022; Chakrabarti et al., 2021). Both branched actin assembly, driven by the Arp2/3 complex and its activators including WASP, and linear filaments assembled by formins are central to different modes of endocytosis (Soykan et al., 2017).

Our screens identified components that drive branched actin assembly, including the Arp2/3 complex (ARPC2, ARPC3, and ARPC4), and the Wiskott–Aldrich syndrome protein (WASP or WAS) (Figure 7A). The WASP- family verprolin homologous protein (WAVE) complex components (WASF2, BRK1, ABI1, CYFIP1) had similar phenotypes, although only NCKAP1L (also known as HEM1) passed our FDR thresholds for significance (Figure 7A). These genes all had modest effects in our integrative analysis for internalization, consistent with findings that branched actin contributes to FPR1 endocytosis (B. C. Subramanian et al., 2018). However, they had stronger effects on basal surface expression, suggesting that branched actin polymerization is important for both FPR1 trafficking to cell surface and its internalization (Figure 5A and 7A).

In contrast to the branched actin components, we identified the formin mDia1 (but not mDia2 or mDia3) as an endocytosis-specific hit (Figure 7A). To validate this finding, we used the formin inhibitor SMIFH2 in dHL-60s and primary human neutrophils. In dHL-60 cells, we detected significant FPR1 internalization defects at SMIFH2 concentrations of 20 μM and higher (Figure 7B). At 100 μM, ∼80% of FPR1 was retained on the cell surface. However, at higher concentrations, SMIFH2 also inhibits myosin motors (Nishimura et al., 2021), and this dramatic internalization defect may be a result of multiple targets affected by the inhibitor. We also observed a decrease in basal surface FPR1 with high SMIFH2 concentrations (Supplementary Figure 6), suggesting a more complex phenotype where different pathways may be affected by the inhibitor treatment. In primary neutrophils, we observed a more pronounced blockage in FPR1 internalization with 50 μM SMIFH2 treatment compared to dHL-60s (Figure 7C). Overall, these results suggest an important role for mDia1 in regulating FPR1 internalization in human neutrophils.

### Arf6 is a key regulator of FPR1 internalization

We also identified Arf6 as an endocytosis-specific hit. Members of the adenosine diphosphate (ADP)-ribosylation factor (ARF) family of small GTPases take roles in membrane trafficking, cytoskeleton remodeling, signal transduction, and regulation of membrane lipid content (Chen et al., 2022; Donaldson & Jackson, 2011). The Arf isoforms expressed in humans have been grouped into 3 classes: class I (ARF1 and ARF3), class II (ARF4 and ARF5), and class III (ARF6). These classes differ in subcellular localization and function. ARF1 GTPase activity regulates trafficking of newly synthesized GPCRs from the Golgi to the plasma membrane (Dong et al., 2010). Consistent with this role, ARF1 was a hit in our FPR1 basal surface expression screen (Figure 8A, Supplementary Figure 7A). Knockout of ARF3, ARF4, or ARF5 did not affect surface FPR1 levels (Figure 8A). However, Arf6, the only plasma membrane-associated Arf (Cavenagh et al., 1996), was a hit specifically for FPR1 internalization (Figure 8A, Supplementary Figure 7A). ARF6 can contribute to both clathrin dependent-endocytosis, through AP2 recruitment (Paleotti et al., 2005; Poupart et al., 2007), and clathrin independent-endocytosis (Delaney et al., 2002), and it has been linked to the endocytosis of other GPCRs (Claing et al., 2001; Delaney et al., 2002; Poupart et al., 2007).

**Figure 8.**
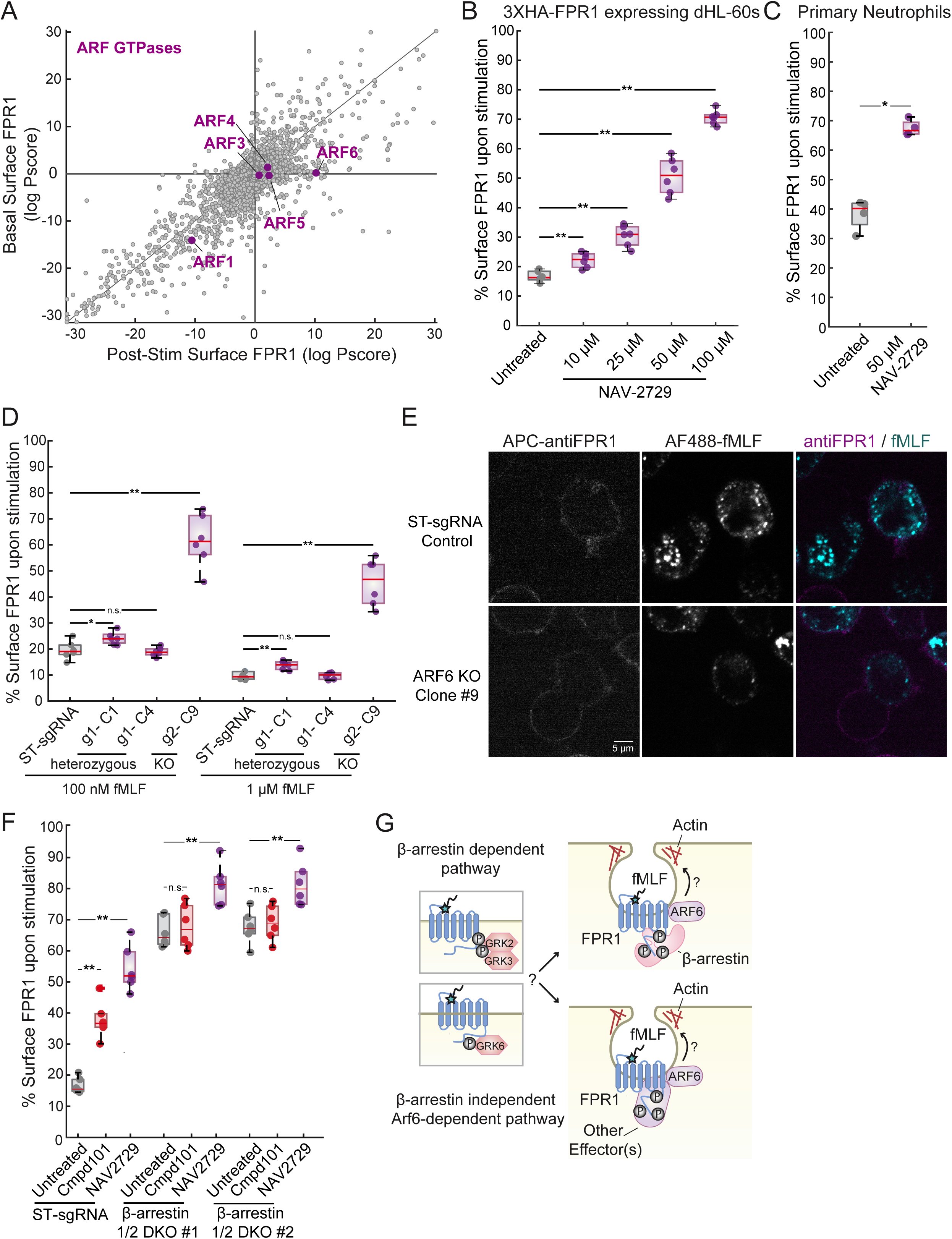
ARF6 is a key mediator of FPR1 internalization. (A) Plot showing the effect of the loss of different ARF genes on FPR1 surface levels in the basal (y-axis) and post-stimulation screens (x-axis). Effect of the Arf6 inhibitor NAV-2729 on FPR1 internalization (B) in dHL-60s (10 to 100 μM) and (C) in primary neutrophils (50 μM). Cells are untreated or pre-treated with NAV- 2729 for 30 mins (n=6 for dHL-60s, n=4 for primary neutrophils). (D) Comparison of FPR1 surface expression in unstimulated and fMLF-stimulated control dHL-60 cells expressing ST sgRNA and clonally selected dHL-60 cell lines expressing ARF6 sgRNA#1 (Clones 1 and 4) or ARF6 sgRNA#2 (Clone 9) (n=6). The mutations (heterozygous or homozygous) were verified using Amplicon Sequencing (see Supplementary Figure 8). (E) ST sgRNA expressing dHL-60s (Control) and ARF6 knockout (ARF6 sgRNA#2 Clone 9) cells were stimulated with AF488-fMLF for 5 min, fixed, and stained with an anti-FPR1 antibody (scale bar, 5μm). (F) ST sgRNA expressing dHL-60s (Control) and β-arrestin double knockout cells were untreated or pretreated with either Compound 101 (10 μM) or NAV-2729 (50 μM) for 30 min and then stimulated with 100 nM fMLF (n=6, n.s. p>0.05, **p<0.01, Mann-Whitney U test). (G) Schematic representation of the potential roles of ARF6 in β-arrestin-dependent and independent endocytic routes used by FPR1.

We used both genetic and chemical approaches to confirm the role of Arf6. First, we used the Arf6 selective inhibitor NAV-2729 in our flow-cytometry based endocytosis assay. We found that the inhibitor causes a strong FPR1 internalization defect in dHL-60s (Figure 8B) and primary human neutrophils (Figure 8C). We note that NAV-2729 can also inhibit the activity of ARF1, and to a lesser extent other regulators of the ARF GTPases (Rosenberg et al., 2023). To further investigate the roles of these two ARF proteins, we generated pooled ARF1 and ARF6 CRISPR knock-out cell lines. In line with our findings from the screens, loss of ARF1 resulted in reduced basal surface FPR1 levels but did not affect its internalization efficiency (Supplementary Figure 7C), while the pooled ARF6 knockout population showed bimodal FPR1 staining after stimulation, where many cells retained FPR1 on their surface (Supplementary Figure 7D).

To understand the heterogeneous behavior of the ARF6 mutant pool, we clonally isolated three cell lines from the pool and characterized them using amplicon sequencing. Two of the clones appeared to be heterozygous, with one functional copy of the ARF6 gene (Supplementary Figure 8A), but the third appeared to be a successful knockout with only mutant sequences (Supplementary Figure 8B). We further confirmed that Clone 9 does not express ARF6 mRNA, while the other clones have similar levels of ARF6 mRNA to safe-targeting sgRNA control cell line (Supplementary Figure 8C). Consistent with their genotypes, the knockout line (Clone 9) retained most of its FPR1 on cell surface after stimulation (Figure 8D, Supplementary Figure 6E), but the heterozygous lines (Clones 1 and 4) could efficiently internalize FPR1 (Figure 8D, Supplementary Figure 8A). We also confirmed the FPR1 internalization defect in the knockout line (Clone 9) using microscopy by the inefficient uptake of the fluorescent ligand (AF488-fMLF) (Figure 8E).

Having established that FPR1 uses both β-arrestin-dependent and -independent internalization pathways, we wanted to investigate whether the role of Arf6 is specific to one of these pathways. We therefore tested whether the Arf6 inhibitor has an additive effect on the FPR1 internalization defect of the β-arrestin DKO cells. For comparison, we treated cells with the GRK2/3 inhibitor in parallel. We found that NAV-2729, but not the GRK2/3 inhibitor Compound 101, resulted in a further increase in FPR1 surface levels after stimulation in these cells (Figure 8F). This finding suggests that Arf6, but not GRK2 or GRK3, participates in β-arrestin independent FPR1 endocytosis (Figure 8G). The moderate effect of the Arf6 inhibitor in the β-arrestin double knockout cell lines suggests that Arf6 may participate in both β-arrestin dependent and independent pathways (Figure 8G).

## Discussion

The levels of chemoattractant receptors at the cell surface can have a major impact on the sensitivity and activity of immune cells, which is relevant for both productive immune responses and inflammatory diseases. Although internalization of many GPCRs in response to stimulation follows well characterized arrestin- and clathrin-mediated pathways, the mechanisms controlling internalization of chemoattractant receptors including FPR1 have been unclear. Previous studies showed a lack of requirement for known mediators of ligand-induced GPCR endocytosis and suggested a non-canonical endocytic route for FPR1. In total, our study indicates that a major challenge for resolving the endocytic mechanism is that multiple pathways likely act in parallel. Using focused experiments, we clarified that β-arrestins and GRKs do play central roles, although no single GRK is required, and the β-arrestin pathway is complemented by a β-arrestin-independent pathway. Moreover, the combined analysis of our genome-wide CRISPR screens provides an overview of protein complexes and components that regulate FPR1 surface expression in different ways, including biogenesis, secretion, internalization and recycling. Future work will be needed to characterize the roles of these mediators in more detail and determine whether some could be therapeutic targets for controlling immune cell function.

### Contributions of GRKs and β-arrestins to FPR1 internalization

The role of GRKs in FPR1 endocytosis was unclear, as GRK2 is a primary mediator of FPR1 phosphorylation (X. Liu et al., 2012; Prossnitz et al., 1995), but it is not required for internalization (B. C. Subramanian et al., 2018). We found that while knockdown of single GRKs does not cause a significant defect, coupling a GRK2/3 inhibitor with GRK6 knockdown strongly impairs FPR1 internalization, suggesting overlapping roles for multiple GRKs (GRK2, GRK3, and GRK6) in FPR1 endocytosis. While GRK2 is well-studied and associated with endocytosis of many GPCRs, GRK6 is less well-studied and its role in FPR1 endocytosis was uncharacterized, even though it is highly expressed in neutrophils. Although our findings suggest that phosphorylation by GRK2 or GRK3 may induce the β-arrestin dependent FPR1 internalization pathway, it is unclear if GRK6 is specifically involved in the β-arrestin independent route. GRK2 and GRK6 can phosphorylate distinct residues on some GPCRs, including the beta-2 adrenergic receptor, leading to different β-arrestin conformations and selective signaling events (Matthees et al., 2021; Nobles et al., 2011). It remains to be uncovered if GRK6 facilitates β-arrestin-independent FPR1 internalization through unique phosphorylation patterns (Liggett, 2011).

Similarly, single knockouts of β-arrestins cause small but significant FPR1 internalization defects, suggesting that β-arrestin1 and 2 are equally important for FPR1 endocytosis. Although amino acid sequences of these two proteins are highly similar (∼80% identical), they differ in their C-terminal region that is critical for regulating β-arrestin binding to GPCRs and other interaction partners such as clathrin (Kee et al., 2024). It is also possible that phosphorylation of FPR1 by GRK2, GRK3 and GRK6 causes binding of different β-arrestins, leading to recruitment of different downstream regulators and signaling effectors, as β-arrestin1 and 2 have unique interaction partners beyond their classic shared effectors (Xiao et al., 2007).

### Regulators of FPR1 trafficking identified through genome-wide CRISPR screens

Our parallel CRISPR knockout screens identified factors controlling trafficking of FPR1 in the basal state and upon stimulation. The identification of genes regulating FPR1 surface expression at different stages may be a valuable resource for studies on trafficking of other chemoattractant receptors or GPCRs. Furthermore, the screening framework and integrative analysis strategy we developed may be applied to future screens to illuminate regulatory mechanisms of receptors in different cell types and experimental conditions.

Our screens revealed involvement of many specific components in FPR1 surface expression and its ligand-induced trafficking. Defects in multiple multi-protein complexes with known roles in receptor recycling and cargo sorting, including Retromer, Retriever, CCC, and HOPS, were found to lower basal FPR1 levels, but also have high FPR1 levels after stimulation, suggesting that these complexes may be involved in both basal secretion of FPR1 and in the internalization process. Besides their roles in cargo recognition and endosomal trafficking, these proteins can interact with an array of effectors for other functions. For instance, COMMD proteins, some of the top endocytosis hits, were reported to interact with a chemokine receptor CXCR4 soon after ligand binding to facilitate GRK6 recruitment (Nakai et al., 2019). A similar involvement of COMMD proteins in FPR1 regulation may explain the strong effect of these mutants in the endocytosis screen, but it has yet to be determined if FPR1 directly interacts with the CCC complex or if CCC complex is involved in GRK or arrestin recruitment to the receptor.

A somewhat surprising result was that defects in the endolysosomal network, such as mutations in WASH complex genes, cause elevated surface FPR1 levels after stimulation. Although receptor internalization has typically been thought to be independent of subsequent endosomal trafficking steps, one potential explanation is that these processes may be linked through feedback. In this case, defects in later stages of endocytosis or the endolysosomal network may cause delays in the earlier steps of internalization. It is also possible that these components are involved in sorting steps that affect surface FPR1 levels by controlling rates of recycling.

### Involvement of formins and Arf6 in FPR1 internalization

To learn about uncharacterized mechanisms of FPR1 internalization, we focused on two endocytosis-specific hits: the formin mDia1 and the small GTPase Arf6. We found that loss of the formin mDia1 or the small GTPase Arf6 is specifically important for the regulation of FPR1 surface levels post-stimulation, suggesting important roles for these proteins in the early steps of FPR1 endocytosis. While mDia1 promotes actin assembly and stabilizes microtubule polymers, Arf6 activity remodels the actin cytoskeleton, controls lipid metabolism, and is implicated in sorting and trafficking of various molecules. Their effects on actin and microtubule dynamics could be important for the formation of tubular invaginations and endosomal vesicles.

Even though there is growing evidence for Arf6’s involvement in actin dynamics and membrane protein trafficking, its specific roles in receptor endocytosis are unclear. GTP-loaded Arf6 is important for the endocytosis and recycling of the beta-2 adrenergic receptor. In this case, Arf6 is recruited to the receptor by β-arrestins, which then leads to a clathrin-dependent endocytic route (Claing et al., 2001). Although Arf6 can also regulate clathrin-independent endocytosis (Donaldson & Jackson, 2011), its direct role in these processes is not established. Arf6 was also a significant hit in genome-wide CRISPR screens for regulators of phagocytosis (Haney et al., 2018) and phagophore closure (Takahashi et al., 2019) in different cell types, suggesting that Arf6 might be a shared component in the vesicle formation step of different biological processes.

We found that Arf6 inhibition reduces FPR1 internalization, even in β-arrestin double knockout cells, suggesting a role for Arf6 in the β-arrestin independent endocytic route. It is possible that Arf6 takes multiple roles in both β-arrestin-dependent and -independent internalization pathways utilized by FPR1, as its inhibition in β-arrestin DKOs did not totally block FPR1 internalization. Consistent with our findings for Arf6, we found that GPCR kinase-interacting protein 2 (GIT2), a GTPase-activating protein (inactivator) for Arf6, was a significant hit, as well. Lack of GIT2 increased basal surface FPR1 levels, but also significantly reduced surface FPR1 following stimulation. This suggests that increased Arf6 activity may result in faster recycling or secretion dynamics in the basal state and a higher internalization capacity upon stimulation.

### Further implications

Mechanistic insights into chemoattractant receptor trafficking could lead to novel strategies to modulate neutrophil behavior in inflammatory diseases. More generally, receptor endocytosis plays a role in a wide range of therapies, including strategies that use receptor internalization to deliver therapeutics into specific cells or to target internalized receptors (Ramírez-García et al., 2019) and therapies based on enhancing surface expression of cell membrane targets (Chew et al., 2020). The mechanisms mediating endocytosis can vary substantially between receptors, and thus characterizing diverse mechanisms of endocytosis is important for developing and improving these therapies.

Contributions of FPR1 endocytosis pathways and their components to neutrophil migration and effector functions require further investigation. Recycling and resensitization of receptors could be critical for maintaining cell surface receptors for sustained migration and signaling. Our previous work indicates that this regulation differs between chemoattractant receptors, in a manner correlated with their signaling behaviors. We found that FPR1 signal attenuation is much slower compared to that of the LTB4 receptor, even though the LTB4 receptor is more resistant to endocytosis (Lundgren et al., 2023). Using multiple endocytosis pathways may allow tuning of FPR1 surface levels, while still rapidly recycling receptors to maintain G-protein signaling.

β-arrestins and other endocytosis regulators may have further contributions to signaling during migration. Indeed, fMLF-dependent localization of β-arrestin1 to the leading edge of migrating neutrophils contributes to directed migration through activation of the small GTPase Rap2 (Gera et al., 2017). Additionally, trafficking of internalized FPR1 is biased towards the leading edge of the migrating neutrophils (B. C. Subramanian et al., 2018). Future studies may uncover whether one or more of the endocytic routes used by FPR1 contribute to this transcytosis of the internalized receptors, and if this process serves as a feedback mechanism to ensure signaling compartmentalization and persistence during cell migration.

In brief, our study addresses discrepancies about the contributions of major endocytic regulators to FPR1 internalization, including presenting evidence for the importance of an overlooked GRK, GRK6. Our CRISPR screens identified many endosome associated multiprotein complexes, trafficking proteins, actin regulators, and signaling pathways that illuminate FPR1 biosynthesis and surface expression in the basal state and its trafficking after chemoattractant stimulation. We validated the roles of two previously unknown facilitators of FPR1 internalization, mDia1 and Arf6, in human neutrophils. Our datasets will inform future studies centered on FPR1, including its role in cell migration. More generally, they should also be relevant for understanding of the regulation of other chemoattractant receptors and GPCRs, including efforts to modulate or co-opt receptor endocytosis for therapeutic approaches.

## Methods

### Cell maintenance and differentiation

All experiments were conducted using differentiated PLB-985 cells, a subline of the HL-60 neutrophil-like cell line. Cells were originally obtained as a gift from O. Weiner (University of California, San Francisco), and their genetic identity was verified by our group before using single-nucleotide polymorphism analysis (Rincón et al., 2018). Cells were grown in RPMI-1640 (Gibco, catalog #72-400-120) media with 9% fetal bovine serum (FBS) (heat-inactivated, Sigma-Aldrich, catalog #F4135 or Avantor, product # 97068-085) and 100 U/ml penicillin-100 mg/ml streptomycin (Gibco, catalog # 15140163) at 37°C and 5% CO2. Cells were maintained at a culture density between 2.0 × 105 and 1.5 × 106 cells/ml. Differentiation into a neutrophil-like state was achieved by growing cells in differentiation medium (RPMI-1640 with 9% FBS, 1.3% Dimethyl sulfoxide (DMSO), and 2% Nutridoma-CS (Roche, catalog #11363743001)) with a starting cell density of 2 × 105 cells/ml. Cell lines were tested for mycoplasma. HEK-293T cells (ATCC CRL-11268) were used for lentiviral production. Cells were cultured in high glucose Dulbecco’s modified Eagle’s medium (Sigma-Aldrich, D5671) that was supplemented with 9% FBS, 1% Glutamax (Gibco, catalog # 35050061), and 100 U/ml penicillin- 100 mg/ml streptomycin. All cell lines were maintained in an incubator at 37°C and 5% CO2.

### Cloning

Exogenous expression of 3XHA-FPR1 is achieved by lentiviral transfection of a EF1a- 3XHA- FPR1-Hygromycin construct in HL-60 cells. 3XHA-FPR1 was amplified in two sequential Gibson assembly reactions. The primers used to generate the construct in this study were given in Supplementary Table 8. These cells were selected using Hygromycin (300 μg/mL) and further purified using the Pierce antiHA-coated magnetic beads (Thermo Scientific, catalog # 88836).

Same backbone vectors were used to clone CRISPR or CRISPRi sgRNA constructs. Backbone vector was digested using BlpI and BstXI FastDigest restriction enzymes (Thermo Fisher), and annealed guide oligos were ligated using Quick Ligase (New England Biolabs, product #M2200S). Guide sequences were given in Supplementary Table 8.

### Stable cell line construction

Stable expression of constructs, 3XHA-FPR1, dCas9, Cas9, CRISPRi and CRISPR sgRNAs, was achieved using 2nd generation lentiviral-mediated gene transfer. The packaging vector used was a gift from Dr. Lifeng Xu (pMD.G). The envelope vector was a gift from Dr. Peter Lewis (pCMV- dR8.2). HEK-293T cells (ATCC CRL-11268) were used for lentiviral production. Cells were cultured in high glucose Dulbecco’s modified Eagle’s medium (Sigma-Aldrich, D5671) that was supplemented with 9% FBS, 1% Glutamax (Gibco, catalog # 35050061), and 100 U/ml penicillin- 100 mg/ml streptomycin. ∼ 5×105 HEK-293T cells were seeded on 6-well TC treated plates. Next day, packaged lentiviral vectors mixed with the transfection reagent PEI in 500 μL Opti-MEM (Gibco, catalog # 51985-034) were added onto cells in 1.5 mL DMEM. After 48 hours, the lentiviral suspension is filtered (PES 0.45µm; catalog # 25-246) and mixed 1-to-1 with HL-60 cells. 24 hours later, infected cells are washed twice with PBS and resuspended in selection media (complete RPMI- 1640 + antibiotic). Centrifugations are done at 200g for 5 min at room temperature.

#### Construction of GRK CRISPRi knockdown cell lines

Lentivirus production for GRK sgRNAs were done in a larger scale using 10 cm plates, and collected lentiviral suspension is concentrated, aliquoted, and flash frozen for multiple use. Single (g1 or g2) CRISPRi knock-downs of GRK2, GRK3, GRK5, and GRK6 were made by stable co-expression of a dCas9 construct (SID4x- dCas9-XTEN80-KRAB) and the guides (ranked #1 and #2 at Weissman Lab hCRISPRi-v2 library, Addgene #1000000090, Supplementary Table 8) through lentiviral infection.

#### β-arrestin, ARF1 and ARF6 CRISPR knockout cell lines

Single (g1 or g2) or double CRISPR knockout cell lines were built by stable co-expression of a Cas9 construct (UCOE-EF1a-Cas9- BFP) and the guides from the Bassik Lab Human CRISPR Deletion Library (Supplementary Table 8). β-arrestin1 sgRNAs were cloned into a vector with a Neomycin marker, and 3xHA-FPR1, Cas9, and β-arrestin2 sgRNA expressing cells were infected with a β-arrestin1 sgRNA to generate the double knockout cell line. Clonal selection is done by limiting dilution in 96-well plates. dCas9 and Cas9 expressing cells were selected with Blasticidin (10 μg/mL), and sgRNA expressing cells were selected using Puromycin (5 μg/mL) or Geneticin (1.5 mg/ml) until the control (non-resistant) cells die for at least 5 days.

To assess the efficiency of Cas9, we used a self-targeting GFP construct (pLV-MLC4-GFP-sgRNA-Puro-T2A-GFP). It was a gift from Michael Bassik and was derived from pMCB306 (Addgene plasmid # 89360). The GFP sgRNA sequence is listed in Supplementary Table 8.

### Neutrophil isolation from blood

Ethical approval for the study of neutrophils from adult healthy controls was granted by the Institutional Review Board from the University of California, Davis (IORG0000251). Participants gave written, informed consent. A microlet lancing device was used to collect blood by finger prick. Neutrophils were isolated using negative selection with EasySep™ Direct Human Neutrophil Isolation Kit (StemCell, catalog #NC0892435), following the manufacturer’s instructions. Immediately after isolation, cells were moved to complete RPMI medium to test effects of different inhibitors in FPR1 endocytosis assays as described below.

### FPR1 internalization assay

2×10^5^ dHL-60 cells in 100 μl were plated on a 96-well Costar assay plate (Corning). 100 μl of either complete medium with DMSO (as a control to measure initial surface FPR1) or 2X fMLF (Sigma-Aldrich, product # F3506) in a complete medium were added using multichannel pipettes simultaneously and mixed well. Cells were incubated at 37°C for either 10 min or 30 min. Samples were then transferred to ice, and centrifugations were performed at 4°C to cease receptor internalization. Two washes were performed after incubation using 150 μl FACS buffer (PBS with 5% heat-inactivated FBS and 0.01% sodium azide). The samples were then stained with 25 μl FACS buffer + APC-antiFPR1 antibody (1:25 dilution, BioLegend catalog # 391610) for 1 hour on ice in the dark. Following immunostaining, two buffer washes are used to remove excess antibody, and cells were resuspended in 200 μl of FACS buffer. Cells were pre-treated with CMPD 101 (Tocris Biosciences, catalog # 5642), NAV-2729 (Sigma-Aldrich Co., SML2238), or SMIFH2 (MedChemExpress, catalog # HY-16931) at 37°C for 30 min where indicated. Surface FPR1 is measured by flow cytometry using a BD FACS Canto II and an LSR II (BD Biosciences, Franklin Lakes, NJ). Unstained cells were used to determine cellular auto-fluorescence (<1%) and subtracted from stained sample measurements. Mean APC signal is used to calculate endocytosis levels with stimulations at different fMLF concentrations or timepoints relative to unstimulated control. Data collected using gates based on FSC and SSC parameters and was analyzed using custom MATLAB scripts. We present data from at least 1×10^5^ live cells for dHL- 60 cells and at least 6×10^4^ cells for primary neutrophil experiments for each condition.

### Sample Preparation and Confocal Imaging

For imaging experiments, 1×10^5^ dHL-60 cells in 100 μl complete medium were plated on a 96- well Costar assay plate (Corning). Cells were pretreated with Compound 101 (Tocris, catalog #5642) where indicated as described above. 100 μl of either complete medium or 2X fMLF- (Cys- AF488)-Amide (Discovery Peptides, catalog # crb1110384h) in a complete medium were added using multichannel pipettes simultaneously and mixed well. Cells were incubated at 37°C for 6 min. Samples were then washed using 150 μl cold FACS buffer to remove unbound or bound but not internalized ligands. Centrifugations were done at 4°C to cease receptor internalization. 100 μl of 4% PFA in PBS was used to fix cells at 37°C for 7 min in dark. Cells were then washed twice with PBS to remove PFA at room temperature. The samples were then stained with 25 μl FACS buffer + APC-antiFPR1 antibody (1:25 dilution, BioLegend catalog # 391610) for 1 hour on ice in the dark. Following immunostaining, two buffer washes are used to remove excess antibody. Fixed cells were plated in 75 μl of a “modified L-15” imaging media (Leibovitz’s L-15 media lacking dye, riboflavin, and folic acid) (UC Davis Biological Media Services) to minimize media autofluorescence on a 96-well glass bottom plate (Celvis, P96-1.5H-N). Images were acquired using a Zeiss Airyscan LSM 800 confocal equipped with a 40x water immersion 1.2- numerical aperture objective and an Axiocam camera. We used ZEN Software settings for AF488 and APC with 488 nm and 640 nm lasers, respectively. ImageJ (Fiji, version 2.14.0) software was used to process images (brightness adjustment, recoloring for presentation, and merging channels).

### Generation of whole-genome CRISPR knockout library of HL-60 cells

Bassik Lab Human CRISPR Deletion Library (Morgens et al., 2017), Addgene catalog #101926- 101934, was packaged into lentiviral particles using TransIT-2020 transfection reagent (VWR, catalog #10767-014). Details about vectors used and HL-60 and HEK-293T cell maintenance were given above. HEK-293T cells were transfected in Optimem medium overnight, and media was changed to complete DMEM the next day. Lentivirus containing supernatant was collected at 24- and 48-hours post media change and filtered (PES 0.45µm; catalog # 25-246). Lentivirus containing supernatant was concentrated 1:50 as described by the manufacturer (Origene, catalog # TR30026). ∼300 million 3XHA-FPR1 expressing HL-60 cells were infected with the concentrated viral particles with sgRNA library in complete RPMI medium containing 2 mg/ml Polybrene (Sigma; catalog # H9268-5G). Following overnight incubation, infected HL-60 cells were washed with PBS. Cells were centrifuged for 10 minutes at 100G and resuspended in the selection medium (complete RPMI-1640 with puromycin). The library was introduced into HL-60 cells with an infection efficiency of ∼30-50% to ensure the representation of all sgRNAs and maximize the number of cells expressing a single sgRNA. The infection efficiency was estimated based on the number of cells that survived infection and showed antibiotic resistance ∼2-days post-infection. Following 5 days of puromycin selection, cells were resuspended in complete RPMI media and maintained at ∼4×10^5^ cells/mL.

### Cell separation and Next-generation sequencing

The pooled libraries were maintained and differentiated at ∼1,000 cells per guide (∼300 million cells) in multiple flasks. Cells differentiated as described above using DMSO and Nutridoma for 6 days. On the day of cell separation, differentiated cells are centrifuged at 100G for 10 min. Cells were resuspended and incubated in either complete RPMI medium (for Surface FPR1 Screen) or complete RPMI with 1 μM fMLF at 37°C for 10 min. Following incubation, cells were washed twice with the FACS buffer. Antibody staining was done using FITC-antiFPR1 (Biolegend, catalog # 391604) on ice for one hour. Top and bottom 30% of the stained cells were sorted on an Astrios EQ (Beckman Coulter) at the University of California Davis Flow Cytometry Shared Resource Laboratory on the day of each screen. Since the sorting step is time consuming, stainings and cell separations were done on separate days for each screen (with less than 24-hour difference), and differentiation of the cells were started accordingly to keep the differences between screens as low as possible except the fMLF stimulation. Genomic DNA was extracted using the QIAamp DNA Blood Maxi Kit (Qiagen, catalog # 51192) according to the manufacturer’s instructions. sgRNA-encoding regions were amplified, barcoded, and Illumina adapters were added in a two-round nested PCR as described in (Morgens et al., 2017) for sequencing. Samples are barcoded using indexed forward PCR primers for identification after pooled sequencing. PCR products (∼280 bp) were purified using double side size selection with SPRIselect beads (Beckman Coulter, product # B23317). Primers used were given in Supplementary Table 8. Sequencing was performed by the UC Davis Genome Center using NextSeq550. Analysis is done using custom MATLAB code that compares enrichment of sgRNAs in low and high bins and integrates data from all four cell populations to pick up hits for different cellular processes (detailed explanation is given in the next section).

### Custom analysis for CRISPR screens

To identify guide RNAs and genes significantly affecting the selection for each of our screens, we used a log-likelihood scoring system that accounts for both the degree of enrichment and the empirical measurement noise. We called the score “CLO” for CRISPR Log Odds. The CLO score uses a negative binomial model to compute the likelihood of observing a specific number of sequencing reads in a sample, after empirically estimating measurement variance as a function of mean read number. We further calculated empirical p-values for each guide and gene, using the distribution of scores for control guide RNAs as an empirical null distribution.

#### Computing CLO scores for individual guide RNAs

We first computed scores for each individual guide RNA as the sign of the effect (positive or negative) times the sum of the log likelihood ratio for obtaining the observed number of reads in each sample, comparing a model in which the true frequency of the guide is equal to the observed frequency against a null model in which the true frequency is the same across all samples.

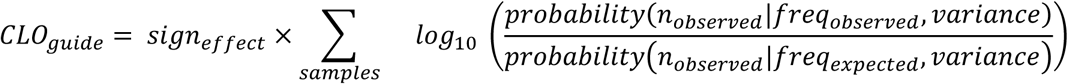

The probability of achieving a specific number of reads in a sample is estimated using the negative binomial model, parametrized by its mean and variance. The mean for a specific guide is equal to the frequency (expected or observed) times the effective total number of reads. The variance is estimated using an empirical strategy described below. For a pairwise comparison of two samples, the sum would be taken over the two samples. However, this strategy can be naturally extended to comparisons involving more samples (e.g., a 4-sample comparison).

Because substantial enrichment or depletion of guides with phenotypes may occur during selection, we use an effective total number of reads, rather than the actual number of reads. We compute the effective number of reads using only the reads for control guides which are expected to have no phenotypic effect. To compute the effective total number of reads for a given sample, we first compute the observed frequency for each control guide, pooling data from all samples to be compared. We then compute a scaling factor as the median (taken over all control guides) of the observed frequency in the sample divided by the frequency in the pooled data. The effective total number of reads for the sample is computed as the actual total reads multiplied by the scaling factor.

Empirical estimates of measurement noise: We estimate the measurement variance for each guide using a variance model, similar to the strategy used by the MAGECK algorithm (Li et al., 2014). However, to avoid over-estimation of variance because of real phenotypic effects of specific guides, we only use the control guides to build the variance models. We model the variance using a linear fit of the log variance as a function of the log mean number of reads. The variance model has two parameters: an intercept and a slope.

For fitting the variance trend for each sample, we compute the deviation values for each guide which are equal to the observed read number minus the expected read number. The expected read number is equal to the pooled frequency (over all samples) times the effective total number of reads. We then pool the deviations for guides with similar expected read numbers (guides are binned by expected read number from 10^1.2^ through 10^3^, with logarithmic step sizes of 10^0.05^). We compute the variance of these deviations, correcting for the number of degrees of freedom in estimating the expected read number (i.e., the variance is multiplied by n_samples_/(n_samples_-1)).

We fit a linear trend to the empirical data for log(variance) as a function of log(expected read number). This provides an estimate of measurement variance for any expected read number value. Finally, if the estimated variance is less than the mean (which would be less than Poisson variability), we apply a minimum bound on the variance equal to 1.01 times the mean.

#### Computing CLO scores at the gene level

After computing guide-level CLO scores, we compute gene-level CLO scores as the sum of the scores for all guide RNAs targeting the gene.

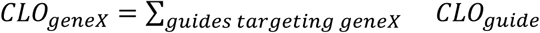

#### Computing p-values for individual guides

As an empirical estimate of the p-value for observing a given CLO score, we use its rank order relative to the scores of all control guides. We define the p-value for hits in the negative direction as 0.5 plus the number of control guides with a lower CLO score, divided by the total number of control guides. We define a p-value for hits in the positive direction as one minus the p-value for the negative direction.

#### Computing p-values at the gene level

We compute p-values at the gene level by combining the guide-level p-values using the algorithm used by (Bailey & Gribskov, 1998). This algorithm computes a single combined p-value based on the null model that the p-values come from a uniform distribution.

#### Computing effect size estimates for individual guides

Raw effect sizes can be computed as the log2 fold-change in frequency of a guide between conditions. While this measure reports on the phenotype associated with the guide RNA, it is heavily influenced by measurement noise, especially when the number of reads for the guide is low for one or more samples. To estimate the effect size given the potential measurement error, we used a Bayesian statistics approach.

The Bayesian approach requires a prior probability distribution for effect size. For the prior distribution, we use a smoothed distribution of the raw effect sizes, considering only guides with an average of at least 100 reads per sample. We smooth the distribution using a standard t-distribution kernel with 3 degrees of freedom. This kernel has a roughly gaussian shape but has long tails which allows effect sizes outside of the range observed among guides with high read counts.

We then use the Bayesian framework to compute posterior likelihoods for the observed read counts in each sample, for possible “true” guide frequencies ranging from the frequency observed in the sample (using the effective total number of reads) to the null expectation (the frequency in the pooled data for all samples), including 20 values at equal intervals in between. We thus compute the likelihoods for a matrix of possible guide frequencies in each sample. For a two-sample comparison, this would be a 21 x 21 matrix of likelihoods. We find the combination of frequencies with the highest posterior likelihood. The estimated effect size is then taken to be the log2 fold-change between these frequencies. These estimated effect sizes range between 0 and the corresponding raw effect sizes.

#### Adjustment to the approach for the 4-sample endocytosis CLO scores

For our endocytosis screen, we compared four samples corresponding to cells with high (top 30%) or low (bottom 30%) surface expression of FPR1 with or without stimulation with the FPR1 ligand fMLF. We expect that guide RNAs affecting cell surface expression of FPR1 generally should have different frequencies in the high versus low samples whether stimulation was applied or not. We also allowed for the possibility that the overall frequency of a guide could be different in the stimulated versus unstimulated samples (perhaps due to random differences in the starting pools of cells).

Finally, we observed empirically that there was a systematic trend relating the degree of enrichment in high versus low samples with stimulation to the degree of enrichment without stimulation. This trend may arise due to the large decrease in surface levels of FPR1 for most cells after stimulation.

To generate CLO scores that most specifically report on phenotypes for endocytosis and recycling of FPR1, we adjust our null model for guide frequencies to allow for differences in guide frequency between high and low samples before stimulation, differences in overall guide abundance in the unstimulated versus stimulated samples, and to correct for the general trend described in the paragraph above. Gene level scores (p-values, signed log p-values, CLO and effect scores) for all genes were given in Supplementary Table 1.

Hits were identified using single tailed multiple hypothesis testing (Storey, 2002) with a positive false discovery rate (pFDR) cut of 0.05. pFDR values were calculated using p-values and the MATLAB function mafdr. Lists for hits identified for different phenotypes were given in Supplementary Tables 2-7.

### Gene Set Enrichment Analysis

Gene set enrichment analysis (GSEA) of ranked gene lists for both screens were performed using the GSEA software (ver 4.3.2) as described by (Reimand et al., 2019). Statistical analysis was performed by the GSEA software using the ranked Log P-scores obtained from our custom analysis described above. For FPR1 internalization, scores from the integrated analysis of the two screens were used. Adjusted p values (FWER) of top 20 gene ontology terms were presented.

### Amplicon Sequencing for Assessing Gene Knockout Efficiency

Genomic DNA was harvested from five million cells for wild-type (WT) and pooled and clonally selected knockout cell lines (Invitrogen Purelink Genomic DNA Kit). The purified gDNA was then used as a PCR template for primers that flank the CRISPR cut site. Primers were given in Supplementary Table 8. The amplified fragments were then PCR cleaned (Zymo DNA clean and concentrator-5) and sequenced using Genewiz Amplicon-EZ service with a NovaSeq 6000 SP in a 2×250bp configuration. The amplicon sequences were analyzed using custom MATLAB codes. Unique sequences that match the primer sequences and longer than 50 bp were identified (>30000 reads/sample). For each remaining unique sequence, the number of reads that identically match the guide sequence were counted. Sequences were aligned to the reference sequence of the gene of interest to determine alterations from the reference sequence. The frequently observed sequences (with read number >1% of the total reads for each sample) were presented.

### Detection of Gene Expression Using Two-Step RT-qPCR

We extracted RNA from differentiated cells on 3 different days. Total RNA extraction is done using Qiagen RNeasy RNA extraction kit according to the manufacturer’s protocol. 1000 ng of purified messenger RNA (mRNA) was used to reverse transcribe to complementary DNA (cDNA) using the 5x All-In-One RT Master Mix kit (abm). Around 20 ng cDNA template is used in the RT- PCR reaction (KiCqStart® SYBR® Green qPCR ReadyMix™) with 3 technical replicates for each condition. 3 rounds of qPCR were performed using cDNA obtained from cells differentiated on different days. Conditions used in the LightCycler® 480 Instrument were: 95°C for 5 s, 45 cycles of 95°C for 10 s, 55°C for 15 s, 72°C for 20 s, Melting curve of 95°C for 5 s, 65°C for 1 m, Cooling. Primers were designed to span exon-exon junctions for GRKs and to amplify a region in Arf6 coding sequence upstream of the sgRNA target sites. Primer sequences are listed in Supplementary Table 8. To quantify enrichment in gene expression in targeting guide expressing cells relative to the safe- or non-targeting guide expressing cells, we used the delta- delta Ct (ΔΔCt) method.

### Statistical analysis

Statistical analysis was performed using MatLab. Comparisons between test and control conditions were performed using the two-sided Mann-Whitney U (Wilcoxon Rank Sum Test) nonparametric test (ranksum function).

## Online supplementary material

The supplementary material provides complementary data to our methodology and findings on the contribution of GRKs, β-arrestins, and Arf6 to FPR1 endocytosis. Fig S1. presents details about the FPR1 internalization assay and key cell lines used in this study. Fig S2 demonstrates qPCR data for each GRK knockdown. Fig S3 provides a panel of line plots for comparison of the FPR1 endocytosis efficiencies in the pooled β-arrestin double knockout cell line and in multiple clones selected from this pool. Fig S4 lists the type of mutations detected in the β-arrestin double knockout pool and clones. Fig S5 demonstrates hits associated with neutrophil differentiation identified in the FPR1 surface expression screen. Fig S6 shows change in surface FPR1 expression at high SMIFH2 concentrations. Fig S7 compares FPR1 surface expression in ARF1 or ARF6 sgRNA expressing cells at the basal and post stimulation level FPR1 expression and FPR1 endocytosis phenotypes. Fig S8 shows the type of Arf6 mutations in the pool and clones and the level of Arf6 mRNA detected in clonal cell lines. Supplementary Table 1 provides all data obtained from the analysis of CRISPR screens. Supplementary Tables 2-7 provides hit lists obtained from the two CRISPR screens and from their integrated analysis. Supplementary Table 8 provides a list of all primers and guide sequences used in this study.

## Data Availability

Sequence data generated from the CRISPR/Cas9 screens and the amplicon sequencing for CRISPR knockout validation are uploaded to the Sequence Read Archive (BioProject ID: PRJNA1246673) and available at http://www.ncbi.nlm.nih.gov/bioproject/1246673. All data is presented in the figures and supplementary material and is openly available as a GitHub project at emelakdogan/FPR1-Endocytosis-Screen-Paper. Materials, analysis tools or custom scripts will be provided by the corresponding authors upon reasonable request.

## Acknowledgments

This work was supported by the NIH under the grants DP2HD094656 and R01GM148769 to SRC, by the UC Davis Dissertation Year Fellowship to EA, and the University of California, Davis. EA and SRC thank Bridget McLaughlin and Jonathan Van Dyke for their technical assistance at the University of California Davis Flow Cytometry Shared Resource Laboratory with funding from the NCI P30 CA093373 (Comprehensive Cancer Center), S10 OD018223, and James B. Pendleton Charitable Trust. RAK acknowledges support from NIH/NCI K99/R00 Pathway to Independence Award CA259218. EA and SRC thanks current Collins lab members and alumni, Prof. Jodi Nunnari, and Assist. Prof. John Albeck for valuable discussions and their input on this project. EA and SRC thank the labs of Jonathan Weissman (Broad Institute) and Neil Hunter (UC Davis) for sharing reagents, protocols, and equipment. EA thanks Abhineet Ram for his contribution to the illustrations.

The authors declare no competing financial interests.

## Author Contributions

EA: Conceptualization, Data curation, Formal analysis, Funding acquisition, Investigation, Methodology, Project administration, Software, Resources, Supervision, Validation, Visualization, Writing – original draft, Writing – review & editing

SML: Investigation, Writing – review & editing

RAK: Supervision, Writing – review & editing

MCB: Supervision, Resources, Writing – review & editing

SRC: Conceptualization, Data curation, Formal analysis, Funding acquisition, Methodology, Project administration, Software, Resources, Supervision, Validation, Visualization, Writing – original draft, Writing – review & editing

## Figure Legends

**Supplementary Figure 1.**
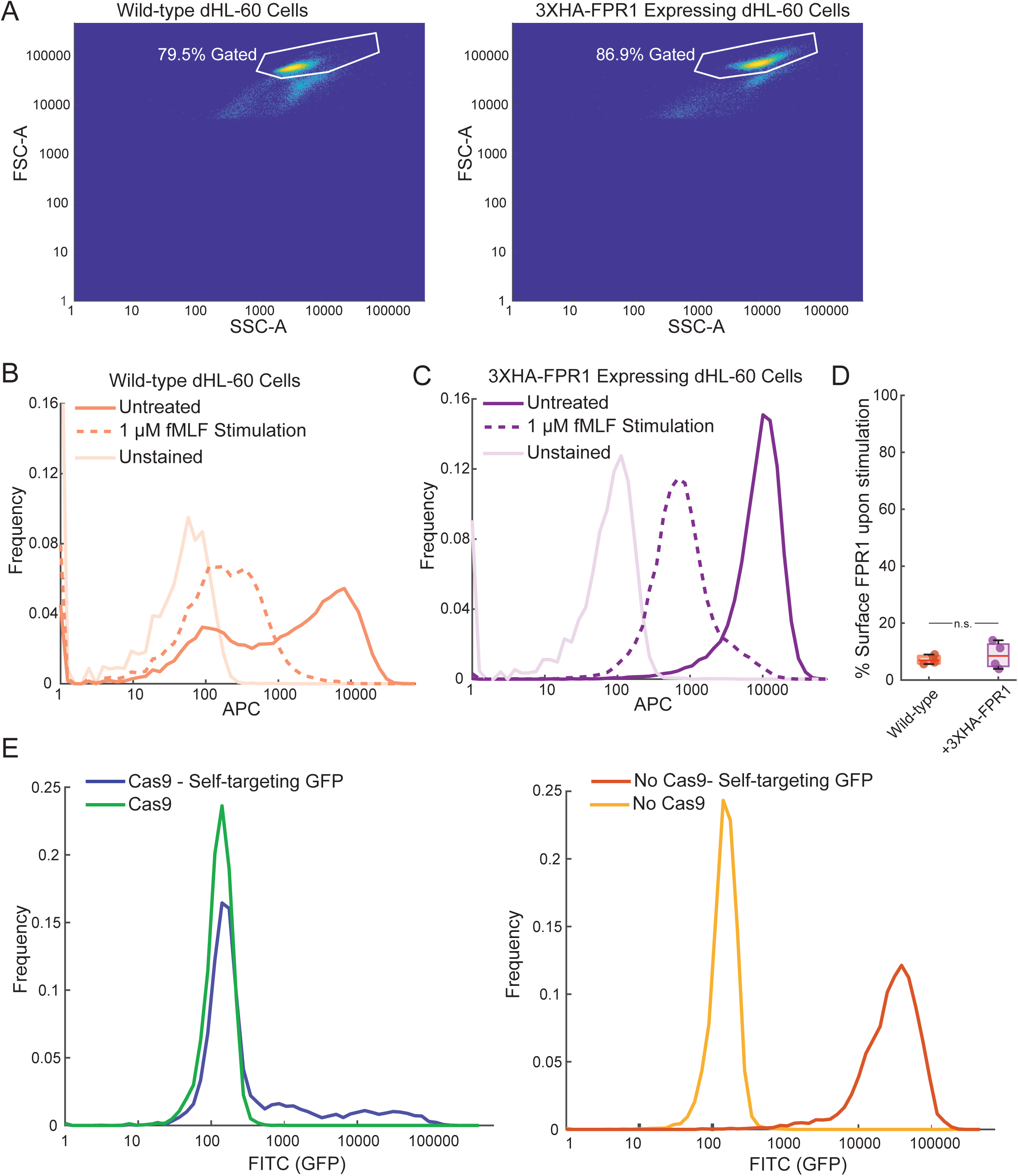
Optimization of tools used in this study. (A) Example gating strategy for flow cytometry measurements. Comparison of FPR1 internalization in (B) wild-type (WT) and (C) 3XHA-FPR1 expressing dHL-60 cells. Cells were untreated or stimulated with 100 nM fMLF and stained for remaining surface FPR1. Note that basal surface expression of FPR1 is much more uniform in cells with the 3XHA-FPR1 construct. (D) FPR1 endocytosis capacities of WT and 3XHA-FPR1 expressing dHL-60s were compared as described in B and C (n=4). Data for wild-type is a subset of the dataset used in Figure 1 and reused in this figure for comparison purposes. (E) Representation of knockout efficiency using a construct that expresses GFP and an sgRNA that targets GFP expression. >80% GFP signal is lost when this construct is expressed along with Cas9.

**Supplementary Figure 2.**
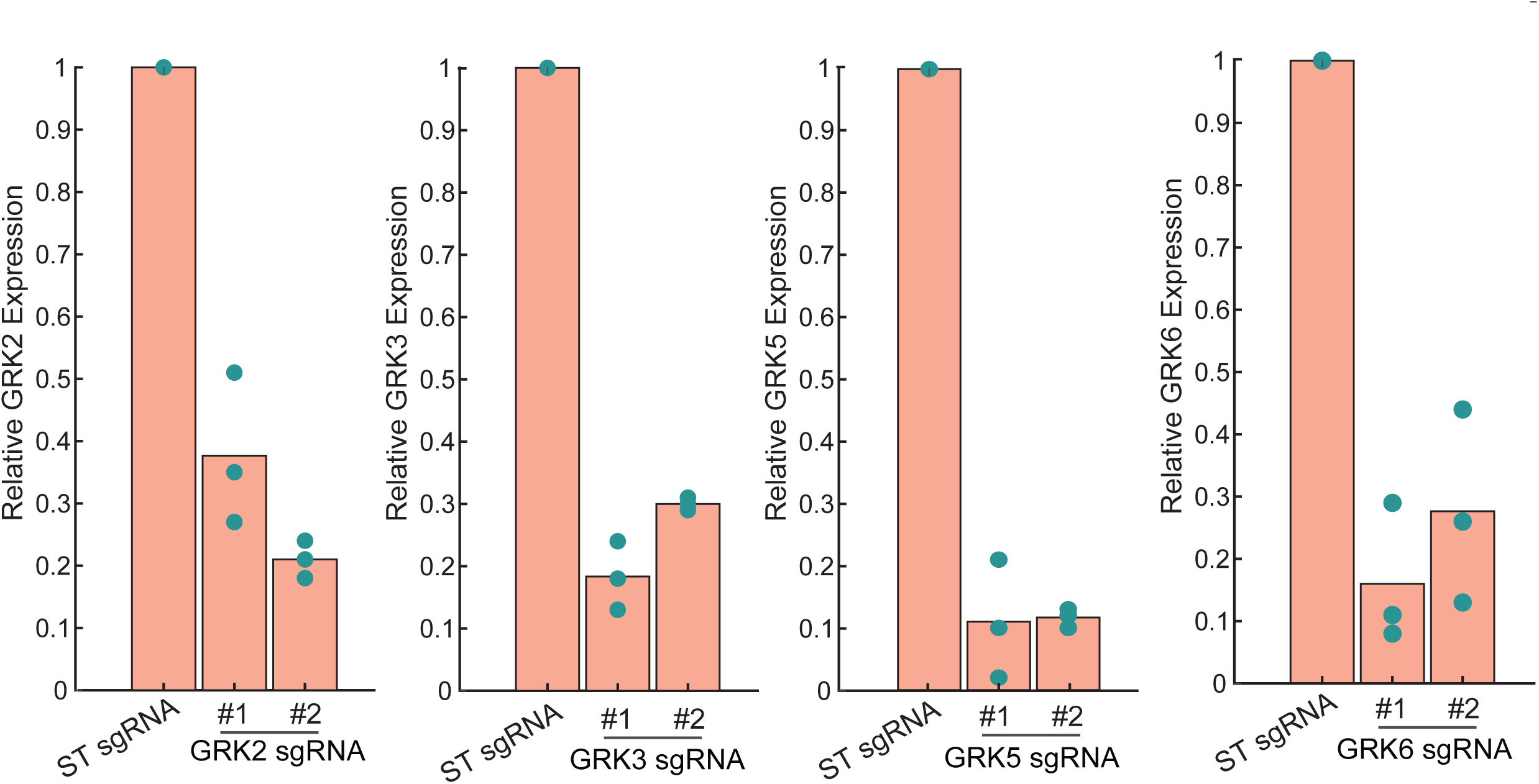
Knockdown efficiencies in dHL-60 cells expressing dCas9 and GRK sgRNAs assessed by RT-qPCR. Dots represent three biological replicates for each condition. Three technical replicates are done in all experiments, and mean Cp values were used for subsequent calculations.

**Supplementary Figure 3.**
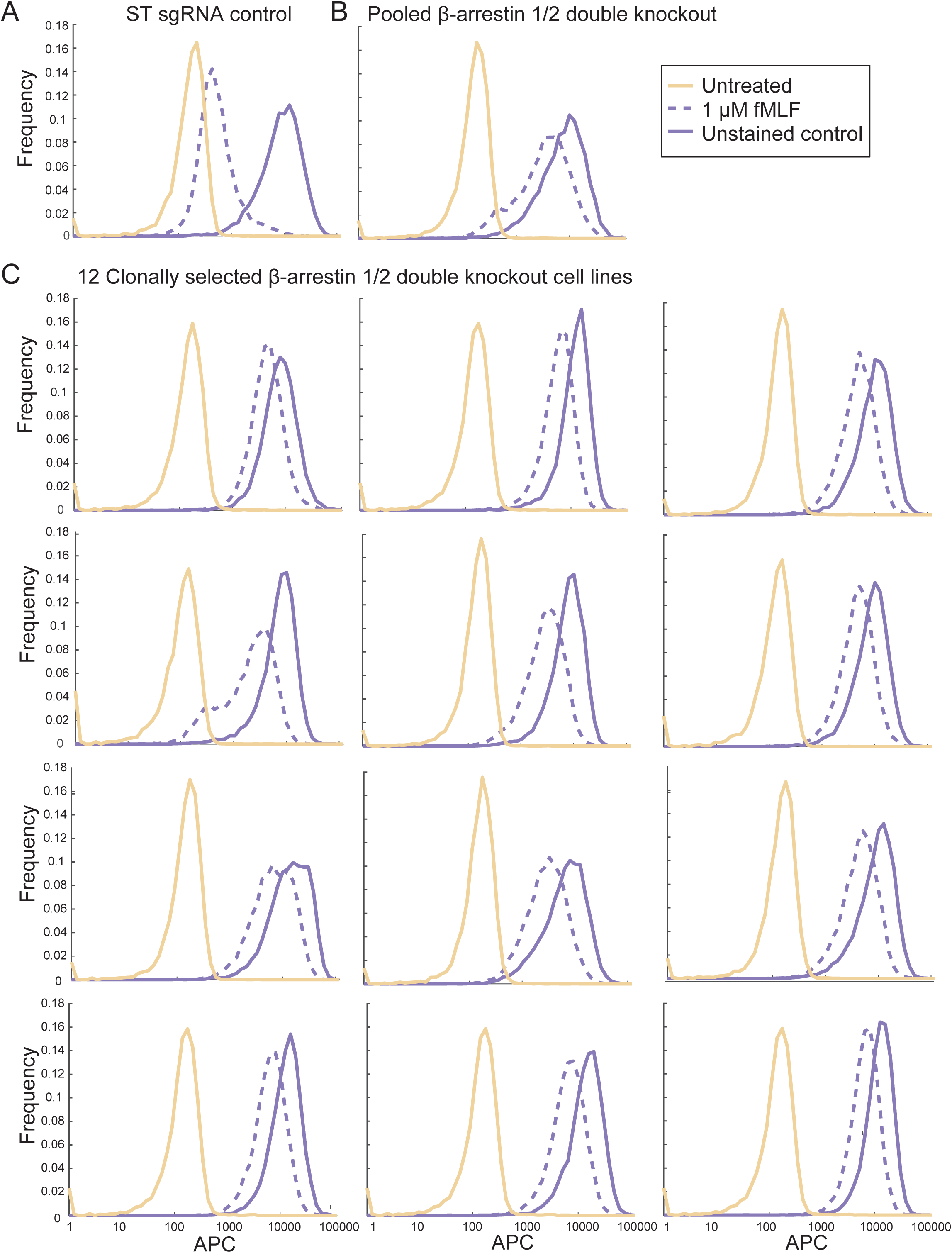
Pooledβ-arrestin DKO has the same endocytosis phenotype as individual clones. Clonally selected β-arrestin double CRISPR knockout cells are compared to pooled double β-arrestin knockout cell lines using the FPR1 endocytosis assay. Cells were untreated or stimulated with 1 μM fMLF. (A) A representative plot showing fMLF-induced decrease in surface FPR1 in the control sgRNA expressing cells. (B) Pooled double β-arrestin knockout cells have a smaller shift after fMLF stimulation. (C) Representative plots obtained from twelve clonally selected β-arrestin double knockout cell lines. Phenotypes observed in clonally selected and pooled knockout cell lines resemble each other.

**Supplementary Figure 4.**
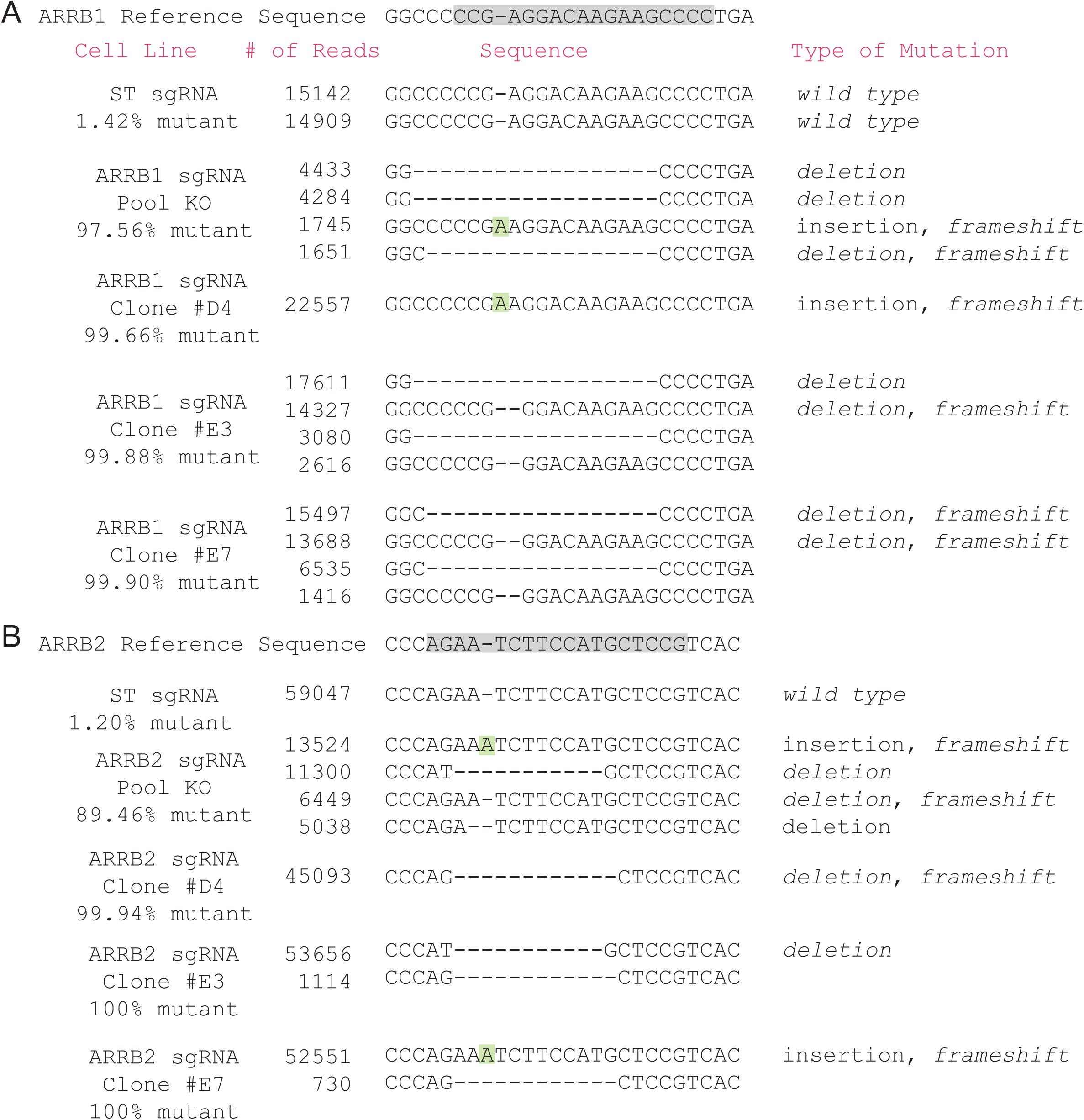
Comparison of amplicon sequencing results forβ-arrestin KO cell lines. Sequences with read numbers greater than 1% of the total reads obtained for the particular sample are presented for (A) β-arrestin1 and (B) β-arrestin2 in control sgRNA expressing cells, pooled and cloned β-arrestin DKO cell lines. Mutant percentage is calculated by subtracting the percentage of reads that contain the expected genomic sequence from 100.

**Supplementary Figure 5.**
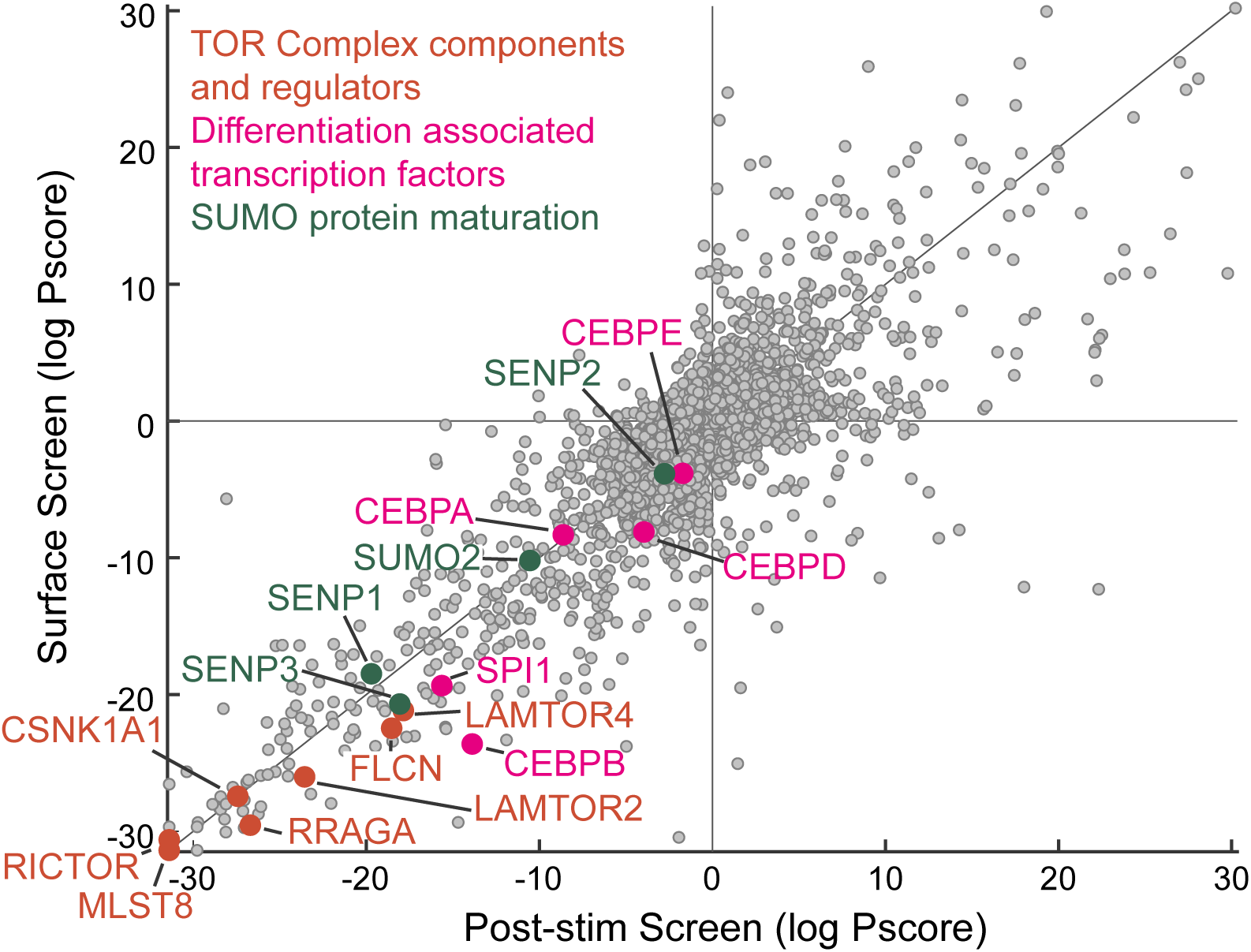
Cell differentiation associated hits identified in the surface FPR1 screen. Plot showing the effect of the knockout of the TOR associated genes, neutrophil differentiation associated transcription factor encoding genes, and genes that function in the maturation of SUMO proteins on FPR1 surface levels in the basal (y-axis) and post-stimulation screens (x-axis).

**Supplementary Figure 6.**
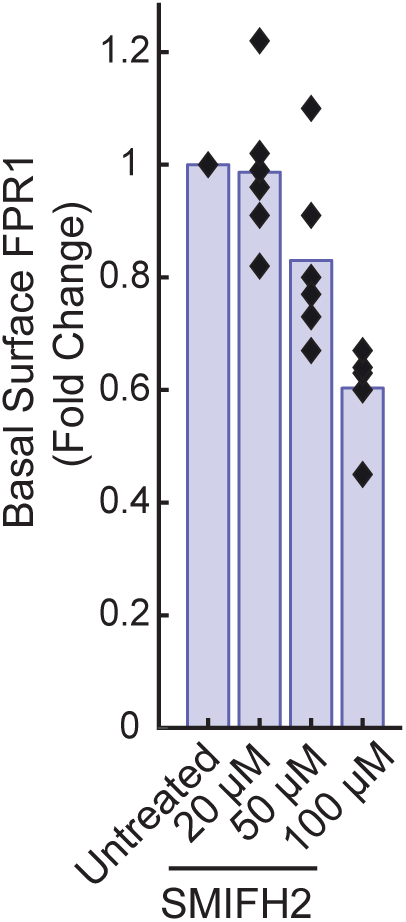
High SMIFH2 concentrations cause decreases in basal surface FPR1. Fold change in surface FPR1 from the control is calculated for different concentrations of SMIFH2 treatment.

**Supplementary Figure 7.**
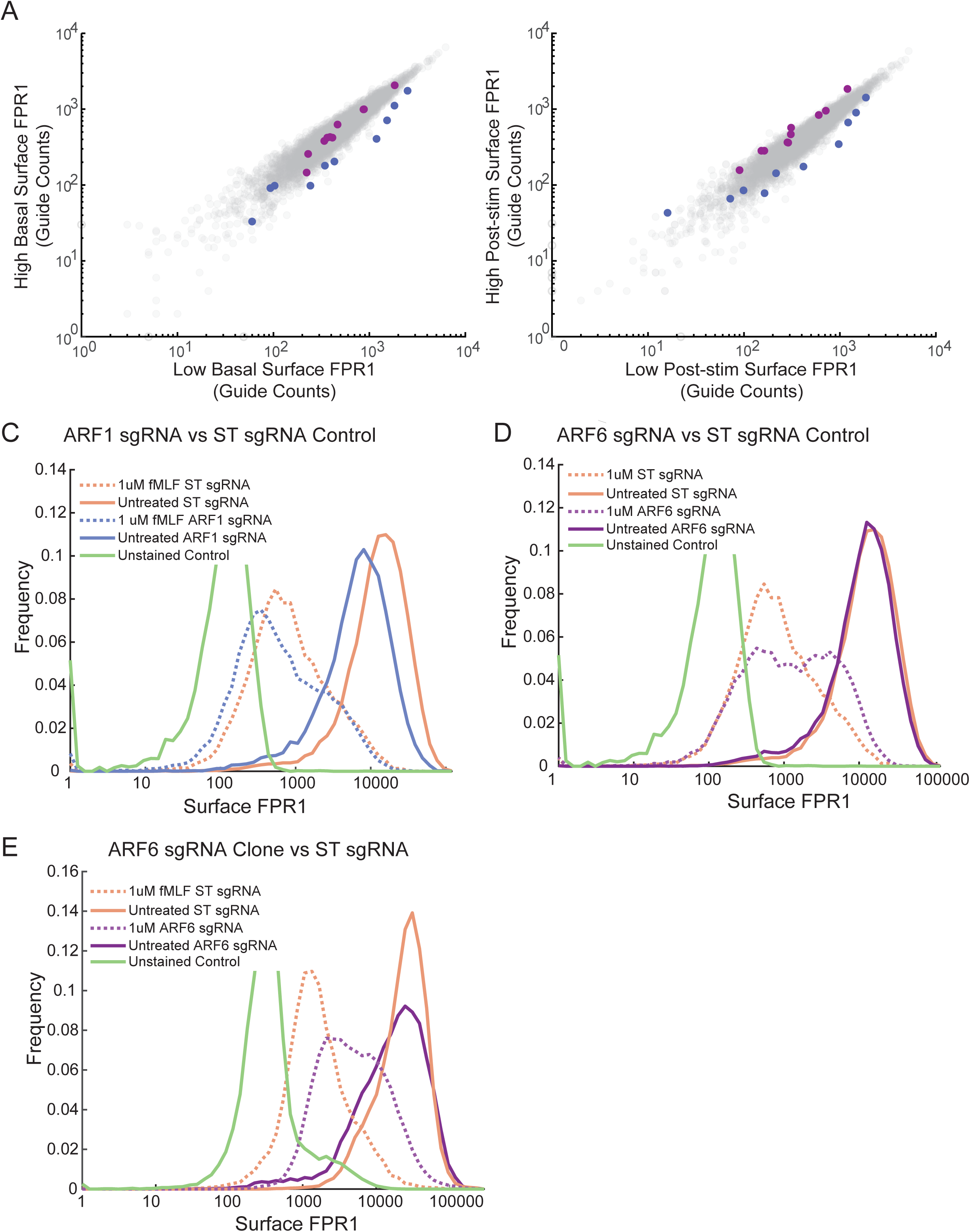
Arf6 facilitates FPR1 internalization, while Arf1 takes roles in basal surface expression. Comparison of read counts for 10 unique sgRNAs targeting ARF1 or ARF6 to the guide counts of the control guides in (A) the surface expression screen and (B) the post-stimulation FPR1 surface expression screen. Shift from the control guides indicates a lower or a higher surface FPR1 in the basal state or after stimulation. Comparison of FPR1 internalization in (C) pooled Arf1, (D) pooled Arf6 (sgRNA#2), and (E) clonally selected Arf6 knockout cells (sgRNA#2, Clone 9) versus ST sgRNA expressing cells.

**Supplementary Figure 8.**
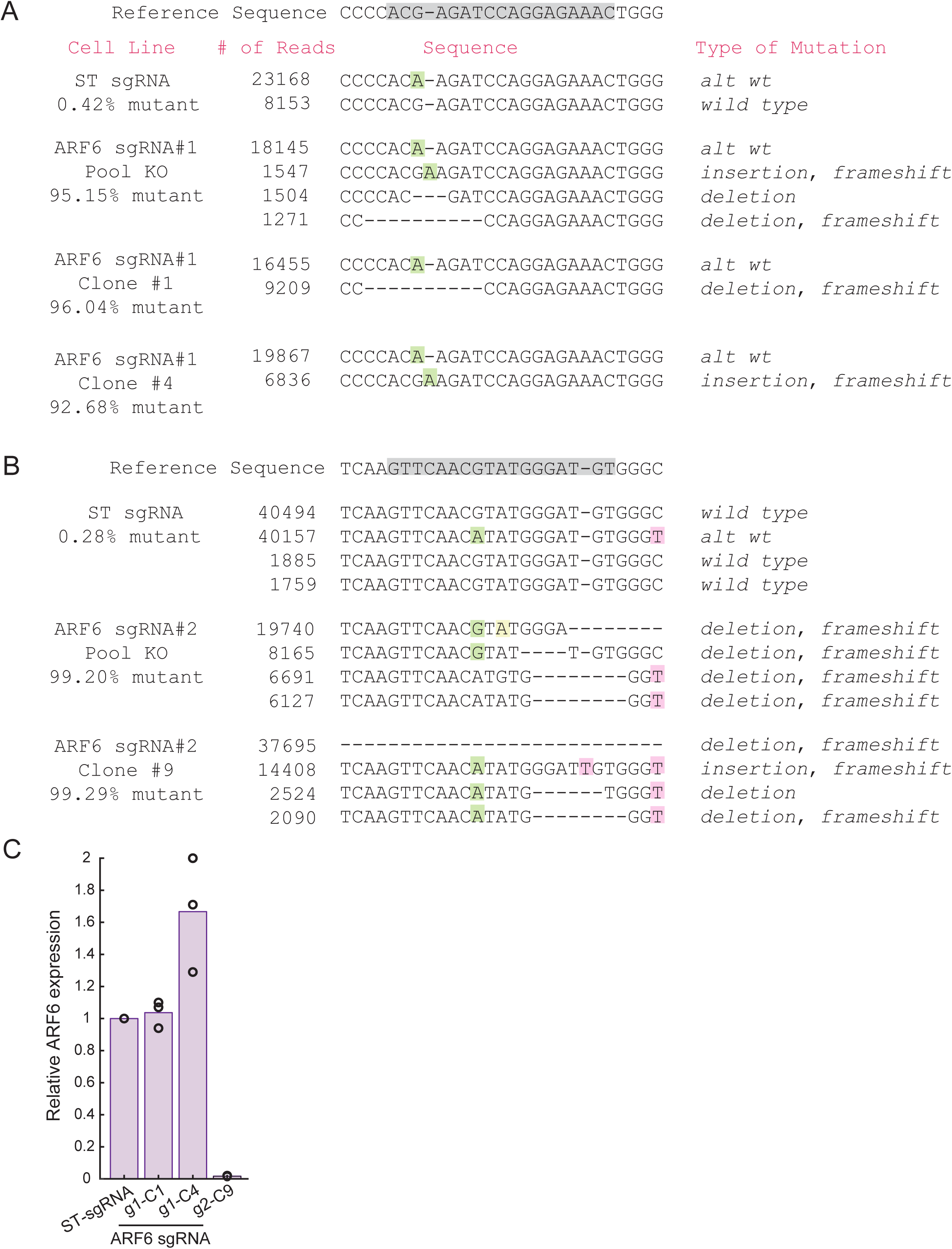
Comparison of amplicon sequencing results for ARF6. Sequences with read numbers greater than 1% of the total reads for each sample are presented. (A) ARF6 sequences for ST sgRNA expressing cells, pooled ARF6 sgRNA#1, and Clones 1 and 4. (B) ARF6 sequences for ST sgRNA expressing cells, pooled ARF6 sgRNA#2, and Clone 9. We detected a frequent base variant in the control cell sequences from the reference sequence (G>A, highlighted in green), and we refer to this variant as the alternative wild-type (alt wt). Mutant percentage is calculated by subtracting the percentage of reads that contain the intact control sequence (wild-type or alt wt) from 100. (C) qPCR to detect presence of ARF6 mRNA in control sgRNA expressing and clonally selected ARF6 sgRNA expressing cells.

**Supplementary Table 1. Screen scores for all genes.**

**Supplementary Table 2. Negative hits identified in the basal surface FPR1 expression screen.**

**Supplementary Table 3. Positive hits identified in the basal surface FPR1 expression screen.**

**Supplementary Table 4. Negative hits identified in the post-stimulation surface FPR1 expression screen.**

**Supplementary Table 5. Positive hits identified in the post-stimulation surface FPR1 expression screen.**

**Supplementary Table 6. Negative hits identified by the integrated analysis of two screens for FPR1 internalization.**

**Supplementary Table 7. Positive hits identified by the integrated analysis of two screens for FPR1 internalization.**

**Supplementary Table 8. Primer and guide sequences used in this study.** Sequences given in 5’ to 3’ direction.

## References

Abouelezz, A., & Almeida-Souza, L. (2022). The mammalian endocytic cytoskeleton. European Journal of Cell Biology, 101(2), 151222. 10.1016/j.ejcb.2022.151222

Bai, S., Hou, W., Yao, Y., Meng, J., Wei, Y., Hu, F., Hu, X., Wu, J., Zhang, N., Xu, R., Tian, F., Wang, B., Liao, H., Du, Y., Fang, H., He, W., Liu, Y., Shen, B., & Du, J. (2022). Exocyst controls exosome biogenesis via Rab11a. Molecular Therapy - Nucleic Acids, 27, 535–546. 10.1016/j.omtn.2021.12.023

Bailey, T. L., & Gribskov, M. (1998). Combining evidence using p-values: Application to sequence homology searches. Bioinformatics, 14(1), 48–54. 10.1093/bioinformatics/14.1.48

Belliveau, N. M., Footer, M. J., Akdoǧan, E., Van Loon, A. P., Collins, S. R., & Theriot, J. A. (2023). Whole-genome screens reveal regulators of differentiation state and context-dependent migration in human neutrophils. Nature Communications, 14(1), 5770. 10.1038/s41467-023-41452-x

Bennett, T. A., Maestas, D. C., & Prossnitz, E. R. (2000). Arrestin Binding to the G Protein-coupled N-Formyl Peptide Receptor Is Regulated by the Conserved “DRY” Sequence. Journal of Biological Chemistry, 275(32), 24590–24594. 10.1074/jbc.C000314200

Bock, C., Datlinger, P., Chardon, F., Coelho, M. A., Dong, M. B., Lawson, K. A., Lu, T., Maroc, L., Norman, T. M., Song, B., Stanley, G., Chen, S., Garnett, M., Li, W., Moffat, J., Qi, L. S., Shapiro, R. S., Shendure, J., Weissman, J. S., & Zhuang, X. (2022). High-content CRISPR screening. Nature Reviews Methods Primers, 2(1), 8. 10.1038/s43586-021-00093-4

Cavenagh, M. M., Whitney, J. A., Carroll, K., Zhang, C., Boman, A. L., Rosenwald, A. G., Mellman, I., & Kahn, R. A. (1996). Intracellular Distribution of Arf Proteins in Mammalian Cells. Journal of Biological Chemistry, 271(36), 21767–21774. 10.1074/jbc.271.36.21767

Chakrabarti, R., Lee, M., & Higgs, H. N. (2021). Multiple roles for actin in secretory and endocytic pathways. Current Biology, 31(10), R603–R618. 10.1016/j.cub.2021.03.038

Chen, P.-W., Gasilina, A., Yadav, M. P., & Randazzo, P. A. (2022). Control of cell signaling by Arf GTPases and their regulators: Focus on links to cancer and other GTPase families. Biochimica et Biophysica Acta (BBA) - Molecular Cell Research, 1869(2), 119171. 10.1016/j.bbamcr.2021.119171

Chew, H. Y., De Lima, P. O., Gonzalez Cruz, J. L., Banushi, B., Echejoh, G., Hu, L., Joseph, S. R., Lum, B., Rae, J., O’Donnell, J. S., Merida De Long, L., Okano, S., King, B., Barry, R., Moi, D., Mazzieri, R., Thomas, R., Souza-Fonseca-Guimaraes, F., Foote, M., … Simpson, F. (2020). Endocytosis Inhibition in Humans to Improve Responses to ADCC-Mediating Antibodies. Cell, 180(5), 895–914.e27. 10.1016/j.cell.2020.02.019

Claing, A., Chen, W., Miller, W. E., Vitale, N., Moss, J., Premont, R. T., & Lefkowitz, R. J. (2001). β-Arrestin-mediated ADP-ribosylation Factor 6 Activation and β2-Adrenergic Receptor Endocytosis. Journal of Biological Chemistry, 276(45), 42509–42513. 10.1074/jbc.M108399200

Conway, J. R., Lex, A., & Gehlenborg, N. (2017). UpSetR: An R package for the visualization of intersecting sets and their properties. Bioinformatics (Oxford, England), 33(18), 2938– 2940. 10.1093/bioinformatics/btx364

Dahlgren, C., Holdfeldt, A., Lind, S., Mårtensson, J., Gabl, M., Björkman, L., Sundqvist, M., & Forsman, H. (2020). Neutrophil Signaling That Challenges Dogmata of G Protein-Coupled Receptor Regulated Functions. ACS Pharmacology & Translational Science, 3(2), 203–220. 10.1021/acsptsci.0c00004

Dahlgren, C., Lind, S., Mårtensson, J., Björkman, L., Wu, Y., Sundqvist, M., & Forsman, H. (2023). G protein coupled pattern recognition receptors expressed in neutrophils: Recognition, activation/modulation, signaling and receptor regulated functions. Immunological Reviews, 314(1), 69–92. 10.1111/imr.13151

Delaney, K. A., Murph, M. M., Brown, L. M., & Radhakrishna, H. (2002). Transfer of M2 Muscarinic Acetylcholine Receptors to Clathrin-derived Early Endosomes following Clathrin-independent Endocytosis. Journal of Biological Chemistry, 277(36), 33439–33446. 10.1074/jbc.M205293200

Donaldson, J. G., & Jackson, C. L. (2011). ARF family G proteins and their regulators: Roles in membrane transport, development and disease. Nature Reviews Molecular Cell Biology, 12(6), 362–375. 10.1038/nrm3117

Dong, C., Zhang, X., Zhou, F., Dou, H., Duvernay, M. T., Zhang, P., & Wu, G. (2010). ADP-Ribosylation Factors Modulate the Cell Surface Transport of G Protein-Coupled Receptors. The Journal of Pharmacology and Experimental Therapeutics, 333(1), 174–183. 10.1124/jpet.109.161489

Dorward, D. A., Lucas, C. D., Chapman, G. B., Haslett, C., Dhaliwal, K., & Rossi, A. G. (2015). The Role of Formylated Peptides and Formyl Peptide Receptor 1 in Governing Neutrophil Function during Acute Inflammation. The American Journal of Pathology, 185(5), 1172– 1184. 10.1016/j.ajpath.2015.01.020

Evers, B., Jastrzebski, K., Heijmans, J. P. M., Grernrum, W., Beijersbergen, R. L., & Bernards, R. (2016). CRISPR knockout screening outperforms shRNA and CRISPRi in identifying essential genes. Nature Biotechnology, 34(6), 631–633. 10.1038/nbt.3536

Gera, N., Swanson, K. D., & Jin, T. (2017). β-Arrestin 1-dependent regulation of Rap2 is required for fMLP-stimulated chemotaxis in neutrophil-like HL-60 cells. Journal of Leukocyte Biology, 101(1), 239–251. 10.1189/jlb.2A1215-572R

Gershlick, D. C., & Lucas, M. (2017). Endosomal Trafficking: Retromer and Retriever Are Relatives in Recycling. Current Biology, 27(22), R1233–R1236. 10.1016/j.cub.2017.10.004

Gilbert, T. L., Bennett, T. A., Maestas, D. C., Cimino, D. F., & Prossnitz, E. R. (2001). Internalization of the Human *N* -Formyl Peptide and C5a Chemoattractant Receptors Occurs via Clathrin-Independent Mechanisms. Biochemistry, 40(12), 3467–3475. 10.1021/bi001320y

Guna, A., Volkmar, N., Christianson, J. C., & Hegde, R. S. (2018). The ER membrane protein complex is a transmembrane domain insertase. Science, 359(6374), 470–473. 10.1126/science.aao3099

Haney, M. S., Bohlen, C. J., Morgens, D. W., Ousey, J. A., Barkal, A. A., Tsui, C. K., Ego, B. K., Levin, R., Kamber, R. A., Collins, H., Tucker, A., Li, A., Vorselen, D., Labitigan, L., Crane, E., Boyle, E., Jiang, L., Chan, J., Rincón, E., … Bassik, M. C. (2018). Identification of phagocytosis regulators using magnetic genome-wide CRISPR screens. Nature Genetics, 50(12), 1716–1727. 10.1038/s41588-018-0254-1

He, H.-Q., & Ye, R. D. (2017). The Formyl Peptide Receptors: Diversity of Ligands and Mechanism for Recognition. Molecules : A Journal of Synthetic Chemistry and Natural Product Chemistry, 22(3), 455. 10.3390/molecules22030455

Herrero-Cervera, A., Soehnlein, O., & Kenne, E. (2022). Neutrophils in chronic inflammatory diseases. Cellular & Molecular Immunology, 19(2), 177–191. 10.1038/s41423-021-00832-3

Hind, L. E., Vincent, W. J. B., & Huttenlocher, A. (2016). Leading from the Back: The Role of the Uropod in Neutrophil Polarization and Migration. Developmental Cell, 38(2), 161–169. 10.1016/j.devcel.2016.06.031

Honda, M., Takeichi, T., Hashimoto, S., Yoshii, D., Isono, K., Hayashida, S., Ohya, Y., Yamamoto, H., Sugawara, Y., & Inomata, Y. (2017). Intravital Imaging of Neutrophil Recruitment Reveals the Efficacy of FPR1 Blockade in Hepatic Ischemia-Reperfusion Injury. The Journal of Immunology, 198(4), 1718–1728. 10.4049/jimmunol.1601773

Kawato, M., Shirakawa, R., Kondo, H., Higashi, T., Ikeda, T., Okawa, K., Fukai, S., Nureki, O., Kita, T., & Horiuchi, H. (2008). Regulation of Platelet Dense Granule Secretion by the Ral GTPase-Exocyst Pathway. Journal of Biological Chemistry, 283(1), 166–174. 10.1074/jbc.M705340200

Kee, T. R., Khan, S. A., Neidhart, M. B., Masters, B. M., Zhao, V. K., Kim, Y. K., McGill Percy, K. C., & Woo, J.-A. A. (2024). The multifaceted functions of β-arrestins and their therapeutic potential in neurodegenerative diseases. Experimental & Molecular Medicine, 56(1), 129–141. 10.1038/s12276-023-01144-4

Kumar, A., Kremer, K. N., Dominguez, D., Tadi, M., & Hedin, K. E. (2011). Gα13 and Rho Mediate Endosomal Trafficking of CXCR4 into Rab11+ Vesicles upon Stromal Cell-Derived Factor-1 Stimulation. The Journal of Immunology, 186(2), 951–958. 10.4049/jimmunol.1002019

Lachance, V., Degrandmaison, J., Marois, S., Robitaille, M., Génier, S., Nadeau, S., Angers, S., & Parent, J.-L. (2013). Ubiquitination and activation of a Rab GTPase promoted by a β2- Adrenergic Receptor/HACE1 complex. Journal of Cell Science, jcs.132944. 10.1242/jcs.132944

Laulumaa, S., & Varjosalo, M. (2021). Commander Complex—A Multifaceted Operator in Intracellular Signaling and Cargo. Cells, 10(12), 3447. 10.3390/cells10123447

Li, W., Xu, H., Xiao, T., Cong, L., Love, M. I., Zhang, F., Irizarry, R. A., Liu, J. S., Brown, M., & Liu, X. S. (2014). MAGeCK enables robust identification of essential genes from genome-scale CRISPR/Cas9 knockout screens. Genome Biology, 15(12), 554. 10.1186/s13059-014-0554-4

Liggett, S. B. (2011). Phosphorylation Barcoding as a Mechanism of Directing GPCR Signaling. Science Signaling, 4(185). 10.1126/scisignal.2002331

Lind, S., Dahlgren, C., Holmdahl, R., Olofsson, P., & Forsman, H. (2021). Functional selective FPR1 signaling in favor of an activation of the neutrophil superoxide generating NOX2 complex. Journal of Leukocyte Biology, 109(6), 1105–1120. 10.1002/JLB.2HI0520-317R

Liu, D., Tsarouhas, V., & Samakovlis, C. (2022). WASH activation controls endosomal recycling and EGFR and Hippo signaling during tumor-suppressive cell competition. Nature Communications, 13(1), 6243. 10.1038/s41467-022-34067-1

Liu, X., Ma, B., Malik, A. B., Tang, H., Yang, T., Sun, B., Wang, G., Minshall, R. D., Li, Y., Zhao, Y., Ye, R. D., & Xu, J. (2012). Bidirectional regulation of neutrophil migration by mitogen-activated protein kinases. Nature Immunology, 13(5), 457–464. 10.1038/ni.2258

Lundgren, S. M., Rocha-Gregg, B. L., Akdoğan, E., Mysore, M. N., Hayes, S., & Collins, S. R. (2023). Signaling dynamics distinguish high- and low-priority neutrophil chemoattractant receptors. Science Signaling, 16(805), eadd1845. 10.1126/scisignal.add1845

Maaty, W. S., Lord, C. I., Gripentrog, J. M., Riesselman, M., Keren-Aviram, G., Liu, T., Dratz, E. A., Bothner, B., & Jesaitis, A. J. (2013). Identification of C-terminal Phosphorylation Sites of N-Formyl Peptide Receptor-1 (FPR1) in Human Blood Neutrophils. Journal of Biological Chemistry, 288(38), 27042–27058. 10.1074/jbc.M113.484113

Matthees, E. S. F., Haider, R. S., Hoffmann, C., & Drube, J. (2021). Differential Regulation of GPCRs—Are GRK Expression Levels the Key? Frontiers in Cell and Developmental Biology, 9, 687489. 10.3389/fcell.2021.687489

Mayer, D., Damberger, F. F., Samarasimhareddy, M., Feldmueller, M., Vuckovic, Z., Flock, T., Bauer, B., Mutt, E., Zosel, F., Allain, F. H. T., Standfuss, J., Schertler, G. F. X., Deupi, X., Sommer, M. E., Hurevich, M., Friedler, A., & Veprintsev, D. B. (2019). Distinct G protein-coupled receptor phosphorylation motifs modulate arrestin affinity and activation and global conformation. Nature Communications, 10(1), 1261. 10.1038/s41467-019-09204-y

Metzemaekers, M., Gouwy, M., & Proost, P. (2020). Neutrophil chemoattractant receptors in health and disease: Double-edged swords. Cellular & Molecular Immunology, 17(5), 433–450. 10.1038/s41423-020-0412-0

Morgens, D. W., Wainberg, M., Boyle, E. A., Ursu, O., Araya, C. L., Tsui, C. K., Haney, M. S., Hess, G. T., Han, K., Jeng, E. E., Li, A., Snyder, M. P., Greenleaf, W. J., Kundaje, A., & Bassik, M. C. (2017). Genome-scale measurement of off-target activity using Cas9 toxicity in high-throughput screens. Nature Communications, 8(1), 15178. 10.1038/ncomms15178

Nakai, A., Fujimoto, J., Miyata, H., Stumm, R., Narazaki, M., Schulz, S., Baba, Y., Kumanogoh, A., & Suzuki, K. (2019). The COMMD3/8 complex determines GRK6 specificity for chemoattractant receptors. Journal of Experimental Medicine, 216(7), 1630–1647. 10.1084/jem.20181494

Nishimura, Y., Shi, S., Zhang, F., Liu, R., Takagi, Y., Bershadsky, A. D., Viasnoff, V., & Sellers, J. R. (2021). The formin inhibitor SMIFH2 inhibits members of the myosin superfamily. Journal of Cell Science, 134(8), jcs253708. 10.1242/jcs.253708

Nobles, K. N., Xiao, K., Ahn, S., Shukla, A. K., Lam, C. M., Rajagopal, S., Strachan, R. T., Huang, T.-Y., Bressler, E. A., Hara, M. R., Shenoy, S. K., Gygi, S. P., & Lefkowitz, R. J. (2011). Distinct Phosphorylation Sites on the β_2_ -Adrenergic Receptor Establish a Barcode That Encodes Differential Functions of β-Arrestin. Science Signaling, 4(185). 10.1126/scisignal.2001707

Novy, B., Dagunts, A., Weishaar, T., Holland, E. E., Adoff, H., Hutchinson, E., De Maria, M., Kampmann, M., Tsvetanova, N. G., & Lobingier, B. T. (2024). An engineered trafficking biosensor reveals a role for DNAJC13 in DOR downregulation. Nature Chemical Biology. 10.1038/s41589-024-01705-2

Paleotti, O., Macia, E., Luton, F., Klein, S., Partisani, M., Chardin, P., Kirchhausen, T., & Franco, M. (2005). The Small G-protein Arf6GTP Recruits the AP-2 Adaptor Complex to Membranes. Journal of Biological Chemistry, 280(22), 21661–21666. 10.1074/jbc.M503099200

Park, S.-J., Nakagawa, T., Kitamura, H., Atsumi, T., Kamon, H., Sawa, S., Kamimura, D., Ueda, N., Iwakura, Y., Ishihara, K., Murakami, M., & Hirano, T. (2004). IL-6 Regulates In Vivo Dendritic Cell Differentiation through STAT3 Activation. The Journal of Immunology, 173(6), 3844–3854. 10.4049/jimmunol.173.6.3844

Peiseler, M., & Kubes, P. (2019). More friend than foe: The emerging role of neutrophils in tissue repair. Journal of Clinical Investigation, 129(7), 2629–2639. 10.1172/JCI124616

Pierce, K. L., Premont, R. T., & Lefkowitz, R. J. (2002). Seven-transmembrane receptors. Nature Reviews Molecular Cell Biology, 3(9), 639–650. 10.1038/nrm908

Potter, R. M., Maestas, D. C., Cimino, D. F., & Prossnitz, E. R. (2006). Regulation of *N* -Formyl Peptide Receptor Signaling and Trafficking by Individual Carboxyl-Terminal Serine and Threonine Residues. The Journal of Immunology, 176(9), 5418–5425. 10.4049/jimmunol.176.9.5418

Poupart, M.-E., Fessart, D., Cotton, M., Laporte, S. A., & Claing, A. (2007). ARF6 regulates angiotensin II type 1 receptor endocytosis by controlling the recruitment of AP-2 and clathrin. Cellular Signalling, 19(11), 2370–2378. 10.1016/j.cellsig.2007.07.015

Prossnitz, E. R., Kim, C. M., Benovic, J. L., & Ye, R. D. (1995). Phosphorylation of the N-Formyl Peptide Receptor Carboxyl Terminus by the G Protein-coupled Receptor Kinase, GRK2. Journal of Biological Chemistry, 270(3), 1130–1137. 10.1074/jbc.270.3.1130

Ramírez-García, P. D., Retamal, J. S., Shenoy, P., Imlach, W., Sykes, M., Truong, N., Constandil, L., Pelissier, T., Nowell, C. J., Khor, S. Y., Layani, L. M., Lumb, C., Poole, D. P., Lieu, T., Stewart, G. D., Mai, Q. N., Jensen, D. D., Latorre, R., Scheff, N. N., … Bunnett, N. W. (2019). A pH-responsive nanoparticle targets the neurokinin 1 receptor in endosomes to prevent chronic pain. Nature Nanotechnology, 14(12), 1150–1159. 10.1038/s41565-019-0568-x

Reimand, J., Isserlin, R., Voisin, V., Kucera, M., Tannus-Lopes, C., Rostamianfar, A., Wadi, L., Meyer, M., Wong, J., Xu, C., Merico, D., & Bader, G. D. (2019). Pathway enrichment analysis and visualization of omics data using g:Profiler, GSEA, Cytoscape and EnrichmentMap. Nature Protocols, 14(2), 482–517. 10.1038/s41596-018-0103-9

Rincón, E., Rocha-Gregg, B. L., & Collins, S. R. (2018). A map of gene expression in neutrophil-like cell lines. BMC Genomics, 19(1), 573. 10.1186/s12864-018-4957-6

Rosenberg, E. M., Jian, X., Soubias, O., Yoon, H.-Y., Yadav, M. P., Hammoudeh, S., Pallikkuth, S., Akpan, I., Chen, P.-W., Maity, T. K., Jenkins, L. M., Yohe, M. E., Byrd, R. A., & Randazzo, P. A. (2023). The small molecule inhibitor NAV-2729 has a complex target profile including multiple ADP-ribosylation factor regulatory proteins. Journal of Biological Chemistry, 299(3), 102992. 10.1016/j.jbc.2023.102992

Sengeløv, H., Boulay, F., Kjeldsen, L., & Borregaard, N. (1994). Subcellular localization and translocation of the receptor for *N* -formylmethionyl-leucyl-phenylalanine in human neutrophils. Biochemical Journal, 299(2), 473–479. 10.1042/bj2990473

Shirakawa, R., & Horiuchi, H. (2015). Ral GTPases: Crucial mediators of exocytosis and tumourigenesis. Journal of Biochemistry, 157(5), 285–299. 10.1093/jb/mvv029

Shurtleff, M. J., Itzhak, D. N., Hussmann, J. A., Schirle Oakdale, N. T., Costa, E. A., Jonikas, M., Weibezahn, J., Popova, K. D., Jan, C. H., Sinitcyn, P., Vembar, S. S., Hernandez, H., Cox, J., Burlingame, A. L., Brodsky, J. L., Frost, A., Borner, G. H., & Weissman, J. S. (2018). The ER membrane protein complex interacts cotranslationally to enable biogenesis of multipass membrane proteins. eLife, 7, e37018. 10.7554/eLife.37018

Soykan, T., Kaempf, N., Sakaba, T., Vollweiter, D., Goerdeler, F., Puchkov, D., Kononenko, N. L., & Haucke, V. (2017). Synaptic Vesicle Endocytosis Occurs on Multiple Timescales and Is Mediated by Formin-Dependent Actin Assembly. Neuron, 93(4), 854–866.e4. 10.1016/j.neuron.2017.02.011

Storey, J. D. (2002). A Direct Approach to False Discovery Rates. Journal of the Royal Statistical Society Series B: Statistical Methodology, 64(3), 479–498. 10.1111/1467-9868.00346

Subramanian, A., Tamayo, P., Mootha, V. K., Mukherjee, S., Ebert, B. L., Gillette, M. A., Paulovich, A., Pomeroy, S. L., Golub, T. R., Lander, E. S., & Mesirov, J. P. (2005). Gene set enrichment analysis: A knowledge-based approach for interpreting genome-wide expression profiles. Proceedings of the National Academy of Sciences, 102(43), 15545– 15550. 10.1073/pnas.0506580102

Subramanian, B. C., Moissoglu, K., & Parent, C. A. (2018). The LTB4–BLT1 axis regulates the polarized trafficking of chemoattractant GPCRs during neutrophil chemotaxis. Journal of Cell Science, 131(18), jcs217422. 10.1242/jcs.217422

Takahashi, Y., Liang, X., Hattori, T., Tang, Z., He, H., Chen, H., Liu, X., Abraham, T., Imamura-Kawasawa, Y., Buchkovich, N. J., Young, M. M., & Wang, H.-G. (2019). VPS37A directs ESCRT recruitment for phagophore closure. Journal of Cell Biology, 218(10), 3336–3354. 10.1083/jcb.201902170

Tiedemann, R. E., Zhu, Y. X., Schmidt, J., Yin, H., Shi, C.-X., Que, Q., Basu, G., Azorsa, D., Perkins, L. M., Braggio, E., Fonseca, R., Bergsagel, P. L., Mousses, S., & Stewart, A. K. (2010). Kinome-wide RNAi studies in human multiple myeloma identify vulnerable kinase targets, including a lymphoid-restricted kinase, GRK6. Blood, 115(8), 1594–1604. 10.1182/blood-2009-09-243980

Vines, C. M., Revankar, C. M., Maestas, D. C., LaRusch, L. L., Cimino, D. F., Kohout, T. A., Lefkowitz, R. J., & Prossnitz, E. R. (2003). N-Formyl Peptide Receptors Internalize but Do Not Recycle in the Absence of Arrestins. Journal of Biological Chemistry, 278(43), 41581– 41584. 10.1074/jbc.C300291200

Wang, J., Chen, M., Li, S., & Ye, R. D. (2019). Targeted Delivery of a Ligand–Drug Conjugate via Formyl Peptide Receptor 1 through Cholesterol-Dependent Endocytosis. Molecular Pharmaceutics, 16(6), 2636–2647. 10.1021/acs.molpharmaceut.9b00188

Wang, J., & Ye, R. D. (2022). Agonist concentration–dependent changes in FPR1 conformation lead to biased signaling for selective activation of phagocyte functions. Proceedings of the National Academy of Sciences, 119(31), e2201249119. 10.1073/pnas.2201249119

Willets, J. M., Challiss, R. A. J., & Nahorski, S. R. (2003). Non-visual GRKs: Are we seeing the whole picture? Trends in Pharmacological Sciences, 24(12), 626–633. 10.1016/j.tips.2003.10.003

Winther, M., Dahlgren, C., & Forsman, H. (2018). Formyl Peptide Receptors in Mice and Men: Similarities and Differences in Recognition of Conventional Ligands and Modulating Lipopeptides. Basic & Clinical Pharmacology & Toxicology, 122(2), 191–198. 10.1111/bcpt.12903

Xiao, K., McClatchy, D. B., Shukla, A. K., Zhao, Y., Chen, M., Shenoy, S. K., Yates, J. R., & Lefkowitz, R. J. (2007). Functional specialization of beta-arrestin interactions revealed by proteomic analysis. Proceedings of the National Academy of Sciences of the United States of America, 104(29), 12011–12016. 10.1073/pnas.0704849104

Xu, J., Wang, F., Van Keymeulen, A., Herzmark, P., Straight, A., Kelly, K., Takuwa, Y., Sugimoto, N., Mitchison, T., & Bourne, H. R. (2003). Divergent Signals and Cytoskeletal Assemblies Regulate Self-Organizing Polarity in Neutrophils. Cell, 114(2), 201–214. 10.1016/S0092-8674(03)00555-5

Zhang, T., De Waard, A. A., Wuhrer, M., & Spaapen, R. M. (2019). The Role of Glycosphingolipids in Immune Cell Functions. Frontiers in Immunology, 10, 90. 10.3389/fimmu.2019.00090

